# Adaptation to postural perturbations under fatigue produces persistent changes in neuromuscular coordination

**DOI:** 10.64898/2026.06.25.734469

**Authors:** Mauro Nardon, Cristiano Alessandro, Tarkeshwar Singh, Matteo Bertucco

## Abstract

Postural control depends on the ability to adapt motor responses to changing environmental and physiological conditions. Neuromuscular fatigue (NMF) is known to alter balance and muscle activation patterns, yet its effects on motor adaptation during whole-body postural tasks and on the persistence of learned strategies remain unclear. This study investigated whether localized NMF of the ankle dorsiflexors influences adaptation to a novel postural perturbation task and whether learning under fatigue induces persistent changes during subsequent re-exposure.

Twenty-five healthy young adults were assigned to either a fatigue or no-fatigue group and completed two experimental sessions separated by 48–72 h allowing recovery from acute fatigue for fatigued group. Participants adapted to repeated mechanical perturbations while standing upright, while ground reaction forces and electromyographic activity of lower-limb muscles were recorded.

NMF did not impair overall adaptation performance, as both groups exhibited similar reductions in performance error across practice. However, participants exposed to fatigue exhibited altered postural recovery dynamics, characterized by a reduced return toward the initial posture following perturbation release. These differences persisted during re-exposure on the subsequent day, despite the absence of acute fatigue. In parallel, NMF modified muscle activation and coactivation patterns involving both fatigued and non-fatigued muscles, several of which were retained during re-exposure.

These findings indicate that the central nervous system preserves successful adaptation to postural perturbations under fatigue by reorganizing neuromuscular coordination and stabilization strategies. Learning under fatigue therefore influences not only immediate motor execution, but also shapes the next-session persistence of postural control strategies.

## INTRODUCTION

Maintaining a standing posture is a non-trivial task that requires continuous active control of postural muscles (Ivanenko and Gurfinkel, 2018). The central nervous system (CNS) employs different strategies to maintain balance depending on environmental conditions, the presence and magnitude of external perturbations, and task-related expectations (Belen’kiĭ et al., 1967; Bouisset and Do, 2008; Nardon et al., 2022). When posture is challenged, postural stability is primarily preserved through the coordinated action of anticipatory postural adjustments (APAs) and compensatory postural adjustments (CPAs). APAs reflect the pre emptive activation of postural muscles based on predictions about an upcoming disturbance, whereas CPAs represent the reactive responses that counteract the perturbation and restore equilibrium once it has occurred (Bouisset and Zattara, 1987; Santos et al., 2010; Cesari et al., 2022). The flexibility and plasticity of these mechanisms enable the motor system to rapidly adapt to changes in task demands or environmental constraints, ensuring efficient and robust postural control (Bertucco and Cesari, 2010; Bertucco et al., 2013, 2021).

Several studies observed that postural control mechanisms and movement execution parameters are impacted by the presence of neuromuscular fatigue (NMF; (Kanekar et al., 2008; Paillard, 2012; Lyu et al., 2021, 2022; Nardon et al., 2024a, 2025)), with physiological and behavioral effects ranging from very short (<1 min) to long-lasting time spans (>1 hour) depending on the exercise duration and modality (Paillard, 2012; Carroll et al., 2017). A recent study observed that the organization and the strength of APA and CPA strategies and their mutual relationship is directly impacted by neuromuscular fatigue (Nardon et al., 2025). Specifically, alterations in muscle activation (both APA and CPA) were more pronounced and prolonged following a localized fatiguing protocol compared to a generalized fatiguing protocol. This aligns with a potential role of fatigue-induced proprioceptive feedback disruption and proprioceptive reweighting (Vuillerme and Boisgontier, 2008; Albanese et al., 2022; Sagnard et al., 2024). There are evidences, however, suggesting that NMF does not impact postural control metrics (Marcolin et al., 2022) and postural control strategies (Remaud et al., 2016). These discrepancies could be attributed to the difference in fatigue-inducing protocols and/or to the targeted muscle group. Despite the observed alterations in muscle activation strategies observed by Nardon et al., participants managed to maintain balance following disturbances, even in presence of acute NMF (Nardon et al., 2025), similarly to the preserved dynamic postural control observed by Marcolin et al. (Marcolin et al., 2022). In our previous study, however, participants had been already exposed to and familiarized with the perturbation (Nardon et al., 2025), limiting insight into how fatigability influences initial adaptation to a novel whole body postural challenge (Krakauer et al., 2019).

When exposed to novel perturbations, the nervous system engages sensorimotor adaptation (Krakauer et al., 2019), a process through which predictive and reactive responses are recalibrated to reduce performance error over repeated exposures. This form of adaptation differs from motor-skill acquisition: whereas skill learning involves developing new movement capabilities, adaptation reflects short-term, error-driven recalibration of existing control policies. Interestingly, evidence from a motor adaptation scenario suggest that postural stability is prioritized over reaching performance while facing a physiological perturbation such as NMF (Nardon et al., 2024b). In fact, reaching performance during a force-field adaptation paradigm worsened for participants exposed to NMF, with muscle activation following differential patterns throughout the adaptation phase in the control and fatigued group. Despite the higher errors in the fatigue group, adaptation curve profiles were maintained, suggesting a delayed – but not totally impaired – adaptation process due to NMF exposure (Nardon et al., 2024b). Similarly, Takahashi et al. observed that internal models formed while exposed to NMF retain some sort of physical effort/muscle activation information, leading to aftereffects in the performance during the re-exposure to the task in a non-fatigued condition (Takahashi et al., 2006).

To our knowledge, only one study has examined long-term consequences of NMF on subsequent re-exposure, showing that learning under fatigue can have detrimental effects that persist even when participants return to practice on the task the following day, without being exposed to NMF (Branscheidt et al., 2019). This study, however, consisted in a very controlled and non-redundant design, where the fatigued muscle was the major contributor to the task. Since findings are inconsistent and mostly limited to short-term effects (Takahashi et al., 2006; Nardon et al., 2024b), it remains unclear whether practicing a novel postural task under fatigability produces persistent next-session changes—that is, differences observable after full recovery, In this study we chose to fatigue the tibialis anterior muscle, which is an important muscle for the control of posture, but is not the sole effector. This allowed us to investigate how the CNS coordinates redundant muscles during adaptation in a fatigued state, and during re-exposure in absence of fatigue. Our previous study (Nardon et al., 2024b) showed that NMF causes an uncoupling between reaching performance and postural stability, with the central nervous system prioritizing stability over reaching accuracy. To prevent such compensatory strategies from masking fatigue effects, we developed a novel task in which postural control is directly tied to task performance, so the two cannot be dissociated.

Therefore, the aims of the study were to investigate how localized NMF of the ankle dorsiflexors influences 1) behavioral adaptation to a novel mechanical postural perturbation and 2) neuromuscular coordination during re exposure 48–72 hours later, when fatigability is no longer present. We predicted that: I) in a multi-joint task such as the control of standing posture, the exposure to NMF would have detrimental effects on the performance, but not on the overall ability to adapt to the perturbation in such a redundant system (i.e., several possible combinations of muscle activation); II) at short term (acute NMF exposure on Day 1), participants in the *Fatigue* group would exhibit differing muscle activation patterns (i.e., lower agonist-antagonist coactivation and greater compensatory activity) both in fatigued and non-fatigued muscles; and III) during the re-exposure to the task in absence of fatigue (Day 2, 48-72 hours after Day 1), differing patterns of muscle activity would persist between groups, as well as a difference in performance.

## MATERIALS AND METHODS

### Participants

Twenty-eight healthy young participants were recruited for the study. Participants had no history of neurological, musculoskeletal disorders, cardiovascular diseases, and did not present conditions or take medications affecting balance. The study protocol was approved by the ethical committee of the University of Verona (approval reference: 2019-UNVRCLE-0298906; date of approval: August 8, 2019) and conformed to the most recent principles of the Declaration of Helsinki. Participants received detailed information about the study before participation and provided written informed consent. Participants were randomly allocated to either a Fatigue (FAT) or No-Fatigue (NoFAT) independently of sex, without stratifying according to sex or anthropometric characteristics. Three participants had to be excluded due to technical problems during the first experimental session (*n*=2) or participant withdrawal from the study following first visit (*n*=1). Demographics of the resulting 25 participants (12 males; 13 females) are reported in **Table 1**.

**Table 1.** Demographics and anthropometric characteristics of the final sample. Values are reported as mean ± SD. Groups did not differ significantly in any of the anthropometric characteristics (independent t-test results: p > 0.05). F: Females, M: Males.

| **Group (F/M)** | **Age (years)** | **Weight (kg)** | **Height (cm)** |
|---|---|---|---|
| Fatigue (5/8) | 21.8 ± 1.6 | 71.4 ± 12.1 | 175.5 ± 9.4 |
| No-Fatigue (8/4) | 22.8 ± 2.7 | 64.4 ± 11.3 | 168.6 ± 9.0 |

### Experimental Design and Set Up

Participants visited the laboratory on two separate occasions (*Day 1*, *Day 2*; 48–72 hours apart), where they were exposed to a paradigm of mechanical perturbation to the standing posture (illustrated in **Fig. 1** and detailed in *Experimental protocol*). Sessions were scheduled either in the morning or in the afternoon and the schedule across sessions was kept the same for each participant. Additionally, to avoid confounders, the time of the day the testing occurred was balanced across groups. Experiments were carried out in the local biomechanics laboratory, equipped with a floor-embedded force plate (model OR-5, AMTI, USA; size: 90 x 90 cm) for the recording of the three components of the ground reaction forces.

**Fig. 1.**
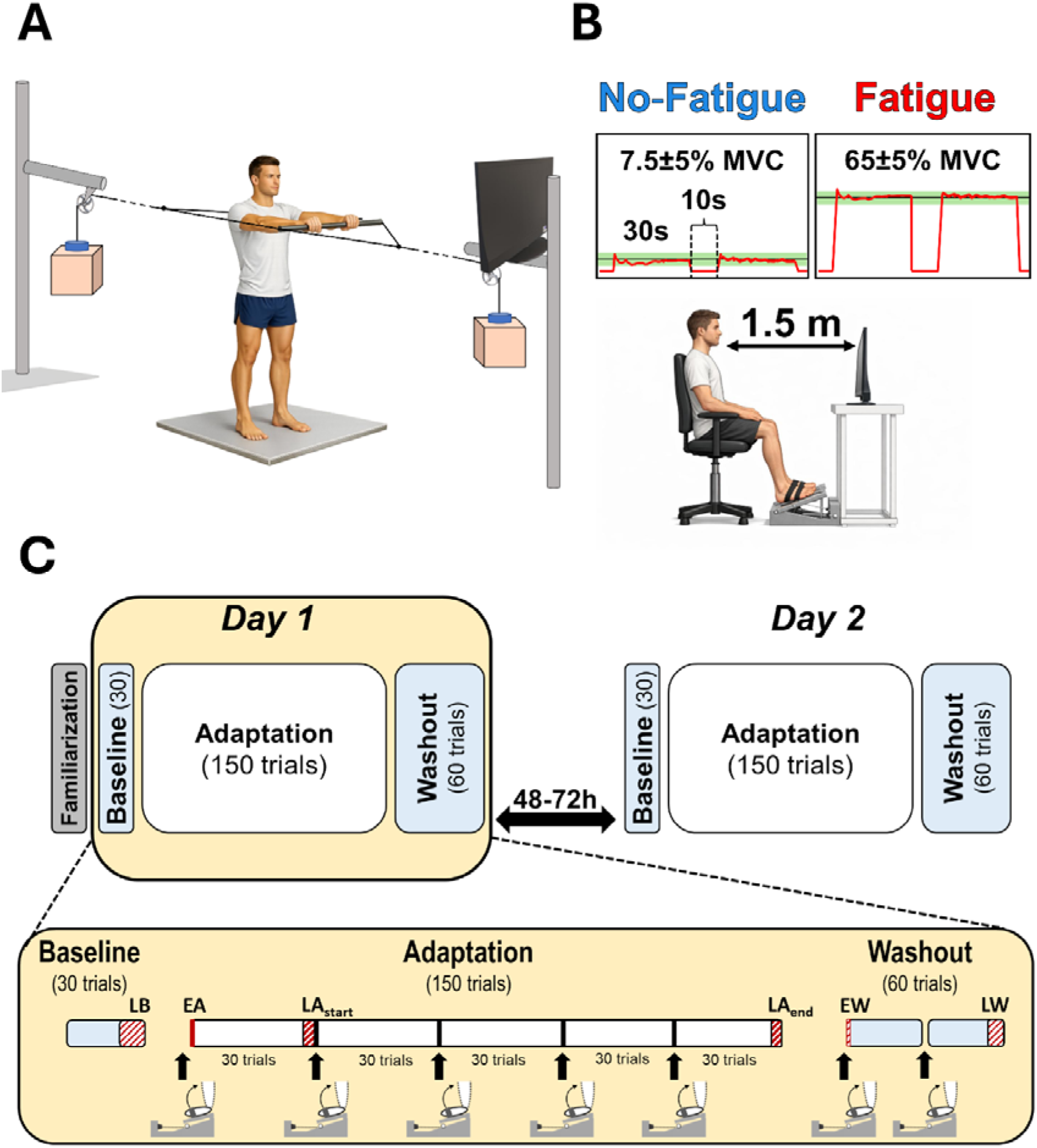
Schematic representation of the experimental setup. A: experimental setup for the postural task. The two boxes in front and at the back of the subject were filled with load plates to match 4% and 8% of participants’ bodyweight to the nearest 50g. Boxes were kept in position by means of electromagnets (shown in light blue) and controlled by an Arduino One board through a button-like switch positioned on the handle. Participants were provided with visual feedback through a monitor in front of them. B: illustration of the fatiguing task (isometric ankle dorsiflexion exercise). Knee and ankle joint angles were kept ∼90° and 110° throughout the exercise. Real-time visual feedback of the force was provided by a monitor screen, which also displayed a shaded area corresponding to the required force target (NoFAT= 7.5 ± 5%; FAT= 65 ± 5% MVC). C: schematic outline of the experimental phases: baseline phase consisted in 30 trials with mechanical load set at 4% of body weight, adaptation phase consisted of 150 perturbed trials (8% of body weight) divided into 5 blocks of 30 trials and washout phase consisted of 2 blocks of 30 trials again at 4% of body weight. Arrows and isometric setup-like symbols at the bottom indicate the fatiguing exercise bouts (see Fatiguing Task in the text). Experimental phases considered for the analyses are highlighted in red. LB: late baseline, EA: early adaptation, LA_start_: late adaptation – start, LA_end_: late adaptation – final block, EW: early washout, LW: late washout.

Surface electromyography (EMG) was recorded from 4 postural muscles: rectus femoris (RF), biceps femoris (BF), tibialis anterior (TA) and soleus (SOL) on the participants’ dominant side using a wireless system (Zerowire, Aurion, Italy). Participants’ skin was properly shaved and cleansed prior to the appliance of Ag/AgCl electrodes (PG10C; Fiab, Vicchio, Italy) with a 20-mm interelectrode distance following recommendations (Hermens et al., 2000). Electrode placement was confirmed by asking the participants to perform a set of isometric contractions and related free movements while observing the resulting EMG patterns (Kendall, 2005). An Arduino One microcontroller (Arduino, Turin, Italy) was used to control power supply of two cylindrical electromagnets keeping two loaded boxes hanging in equilibrium (**Fig. 1A**), one in front- and one in the back of the participant). A button-shaped switch released the box in front of the participant when pressed. The switch was secured to a custom-built aluminum handlebar and positioned in a comfortable position to be pressed with the index of the dominant hand. On switch activation, a square-like signal generated by the Arduino board was also used to trigger the data acquisition system (hardware: MX Control, VICON, Oxfordshire, UK) and align the signals. Data capture was synchronized in Vicon Nexus software (Nexus 2, version 2.12, Vicon Motion Systems Ltd, UK) and set to record for 4 seconds: 2 seconds before and 2 after the trigger pulse. Ground reaction forces and EMG data were sampled at 1000 Hz and synchronized using a hardware device (MX Control, VICON, Oxfordshire, UK). Data were later exported to Matlab software (R2023a, version 9.14.0, MathWorks, Natick, MA, USA) for offline analysis.

A customized frame equipped with a uniaxial strain-gauge force sensor (Model S-AL-A, Deltatech, Sogliano al Rubicone, Italy) was used to perform the localized isometric fatiguing task (**Fig. 1B**; detailed in the dedicated paragraph *Fatiguing Task*). Force data were collected using a computer-based data acquisition and analysis system (hardware: PowerLab 16/30; ML880, ADInstruments, Colorado Springs, CO; software: LabChart 8, ADInstruments, Colorado Springs, CO) at 1000 Hz sampling rate.

### Experimental protocol

During the first visit (*Day 1*), participants received instructions about the aim of the postural task: “minimize the perturbation-induced displacement of your center of pressure (CoP)”. i.e., “try not to be destabilized by the release of the load”. Furthermore, based on pilot tests and due to the novelty of the task, 30 familiarization trials were granted before starting the first session to experience the postural perturbation and stabilize their performance. The entire experimental session lasted ∼3 hours on each visit. Visits were scheduled 48–72-hour apart to avoid accumulation and/or long-lasting effects of fatigue between sessions.

#### Postural Task

Participants were asked to stand barefoot at the center of the force plate with their knees slightly bent and feet in parallel at hip width. Feet position was checked and marked on the surface of the force plate to ensure consistency across experimental blocks. A 22-inch monitor positioned ∼1.8 m in front of the participants was used to provide visual feedback. Participants held the horizontal handlebar at mid-chest level, with their elbows fully extended. The handle was connected to an adjustable system consisting of iron cables and pulleys, with the cylindrical electromagnets connected on both ends that ensured the attachment of two mechanical loads: one in front of the participant and one behind the participant’s body (**Fig. 1A**). The content of the boxes was masked by a tissue cover, preventing participants from visualizing the contained load plates. Participants, by pressing on the switch button positioned on the handlebar, turned off the electromagnet in front of them, causing the release of the load and leading to a mechanical perturbation in the opposite direction (i.e., the handlebar was pulled backwards). Both loads were equivalent in weight, having no effects on posture when the system was in balance. Height of the structure was adjusted to each participant’s height to have the pulley system acting at mid-chest level and to have the feedback monitor at participant’s eye level. The mechanical perturbation paradigm was developed based on a modified version of the setup used in a recent study (Piscitelli et al., 2017).

A calibration trial was conducted at the very beginning of each session to determine the average CoP position in the antero-posterior direction (CoP_AP_) during natural standing. Participants were asked to stand and fixate a point on the screen; hands were positioned on the handle with arms fully extended and parallel to the ground. CoP_AP_ position was recorded for 5 seconds, visually inspected offline to exclude abnormal spikes in CoP position and then averaged for the entire time window. This value was then saved and used 1) to set the zero target (CoP_Target_) for online visual feedback in the successive postural task and 2) to normalize CoP position data during offline analysis of the trials.

During the postural task, participants were initially prompted by the experimenter to assume the starting position (hands on the handle, arms fully extended, knees slightly bent; **Fig. 1A**). CoP_Target_ was projected on the screen as a thin white horizontal line enclosed by a ±2 cm green-shaded area. Real-time CoP_AP_ position was automatically computed by a customized script in MATLAB (R2018a, version 9.4; MathWorks Inc., MA, Natick, USA) which was developed to access real-time data through Nexus SDK files and was superimposed on the screen by means of a thick, horizontal blue line. Participants were instructed to maintain their CoP_AP_ within the green-shaded area. After a “ready” cue, the feedback was turned off following a random delay and participants were required to press as fast as possible the switch that released the load. Visual representation of participant’s CoP_AP_ displacement from CoP_Target_ was provided right after the trial termination (CoP_feedback_) through an arrow superimposed on the CoP_Target_ and its relative value in cm and sign (positive if backward, negative if forward respect to the CoP_Target_). If the error was within the tolerance (COP_Target_ ±2 cm; green-shaded area), the arrow and its value were shown in black color, otherwise the color was in red. Recorded data encompassed a 4-s window, centrally aligned on the instant the switch was pressed (*t_0_*; i.e., 2 seconds before/after pressing the button). Once the trial ended and the file was saved, offline CoP_AP_ displacement computation was performed (CoP_feedback_ = CoP_AP_ – CoP_Target_). Average trial duration was ∼10 seconds.

During *familiarization*, *baseline* and *washout* trials the load plates in the boxes were set to match (to the nearest 50 grams) the 4% of participant’s body weight. Following the *baseline* block, each successive block was preceded by a bout of sustained isometric exercise (details in *Fatiguing Task*). During the trials in the five successive blocks (*adaptation*), another set of boxes, prepared with an 8% body weight load, were used. Participants were not aware of this change. During the final two blocks (*washout*) the boxes were switched back to the initial load (4% of body weight). The choice of loads in percentage to body weight was done to control the amount of perturbation and scale it to anthropometric differences among participants. Percentage values are chosen to produce a consistent, yet controllable perturbation, based on pilot tests where multiple combinations were tested. Sessions were identical in the structure of the protocol (*baseline*, *adaptation*, *washout*), except for familiarization, which was performed only on Day 1. Furthermore, participants in both groups were not exposed to fatigue on Day 2. An outline of the experimental protocol is shown in **Fig. 1C**.

#### Fatiguing Task

Participants were seated in front of a screen (∼1.5 meters away) with their feet fastened to a customized metal frame through two inextensible straps, placed proximally to the metatarsophalangeal joints (See **Fig. 1B**). Knee and ankle joint angles were maintained at approximately 90° and 110°, respectively and the position did not change throughout the entire experimental session. The isometric fatiguing protocol was modulated by previously recorded maximum voluntary contraction (MVC; 3 repetitions of sustained 5 s contractions) of ankle dorsiflexors, and consisted in performing 40s sustained isometric contractions, duty-cycle of 75% and intensity of 65 ± 5% MVC (No-Fatigue: 7.5 ± 5% MVC). Individual baseline MVCs were repeated on each session to account for day-to-day force fluctuations (Dutra et al., 2023) and to confirm effective recovery in the FAT group. In previous studies this protocol produced a significant sustained reduction in participants’ ankle dorsiflexors MVC force, lasting more than 10 minutes. The fatiguing exercise was repeated multiple times (7 bouts: one before each experimental block – except for baseline), and duration of each block of postural task was limited to 10 minutes, to minimize the likelihood of substantial recovery throughout the experiment (Reid et al., 1993; Birtles et al., 2002; Thompson et al., 2015; Carroll et al., 2017).

In the present study, the subjective perception of fatigue was confirmed by the reporting of a Borg RPE (1-10) score of >8 at the end of each exercise bout. A detailed description of the fatiguing protocol has been reported elsewhere (Ricotta et al., 2023; Nardon et al., 2024b).

Exercise time for each bout of isometric tasks of the FAT group was averaged across subjects and rounded up to the nearest cycle (40s). These exercise times were later used to stop participants in the NoFAT group (exercising around 7.5% MVC). This was done to ensure a similar time duration in both groups and required the FAT group to be tested before NoFAT. Participants in both groups were stopped by the experimenter and were blind to exercise intensity (%MVC), termination criteria and exercise bouts duration. The setup for the fatiguing task was positioned ∼3 meters away from the force plate to ensure a rapid transition between the tasks.

### Data analysis

Data was exported from Nexus software and pre-processed offline in MATLAB (Math-Works Inc., Natick, MA, USA, R2025b). Ground reaction forces signals (and CoP) were filtered with a 15 Hz low-pass, 6^th^ order, zero-phase digital Butterworth filter. EMG signals were detrended, band-pass filtered with a 20–450 Hz, 4^th^ order, zero-phase digital Butterworth filter and rectified. Trigger signal from Arduino One was low-pass filtered (10 Hz, 2^nd^ order Butterworth) to confirm alignment of the data to the instant the button was pressed (i.e., the release of the load; *t_0_*). Aligned and pre-processed datasets were stored in matrices for further analysis.

Adaptation and washout processes were evaluated – separately – by comparing variables at six stages of the experiment on each session (Fig. 1): late baseline (LB; last 10 trials in baseline phase); early adaptation (1^st^ trial in adaptation phase); late adaptation – start (last 3 trials of the 1^st^ adaptation block); late adaptation – final block (last 5 trials in adaptation phase); early washout (1^st^ trial in washout phase); late washout (last 5 force trials in washout phase). Different numbers of trials were used across phases to capture phase-specific adaptation dynamics, in accordance with previous similar studies (Ahmed and Wolpert, 2009; Pienciak-Siewert et al., 2020; Nardon et al., 2024b). A schematic outline of the experimental phases and the relative windows used for the analysis are shown in **Fig. 1C**.

#### CoP variables

Performance error (PE) was computed from CoP_AP_ signal – similarly to the online feedback provided to the participants at the end of each task – as the peak backward (i.e., positive) displacement of the CoP from the target position (CoP_Target_) over a 1s-time window after *t0* (**Fig. 2A**). Additionally, we compared adaptation profiles and adaptation rates of the two groups by considering PE values during the first block of trials during the adaptation phase (n = 30 trials), in line with similar approaches in previous studies (Takahashi et al., 2006; Malone et al., 2011). To reduce trial-to-trial variability and lower the influence of trials largely deviating from the trends, while preserving participant-specific adaptation profiles, individual data were smoothed using a 3-point moving average filter. A single-term exponential function was then fitted to each participant’s data in MATLAB using the following formula:

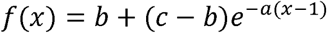

where *a* represents the exponential decay rate, *b* the asymptotic value, and *c* the estimated PE at the first adaptation trial. Fitting parameters were then compared between groups to assess potential differences in adaptation profiles and rates on each day separately.

**Fig. 2.**
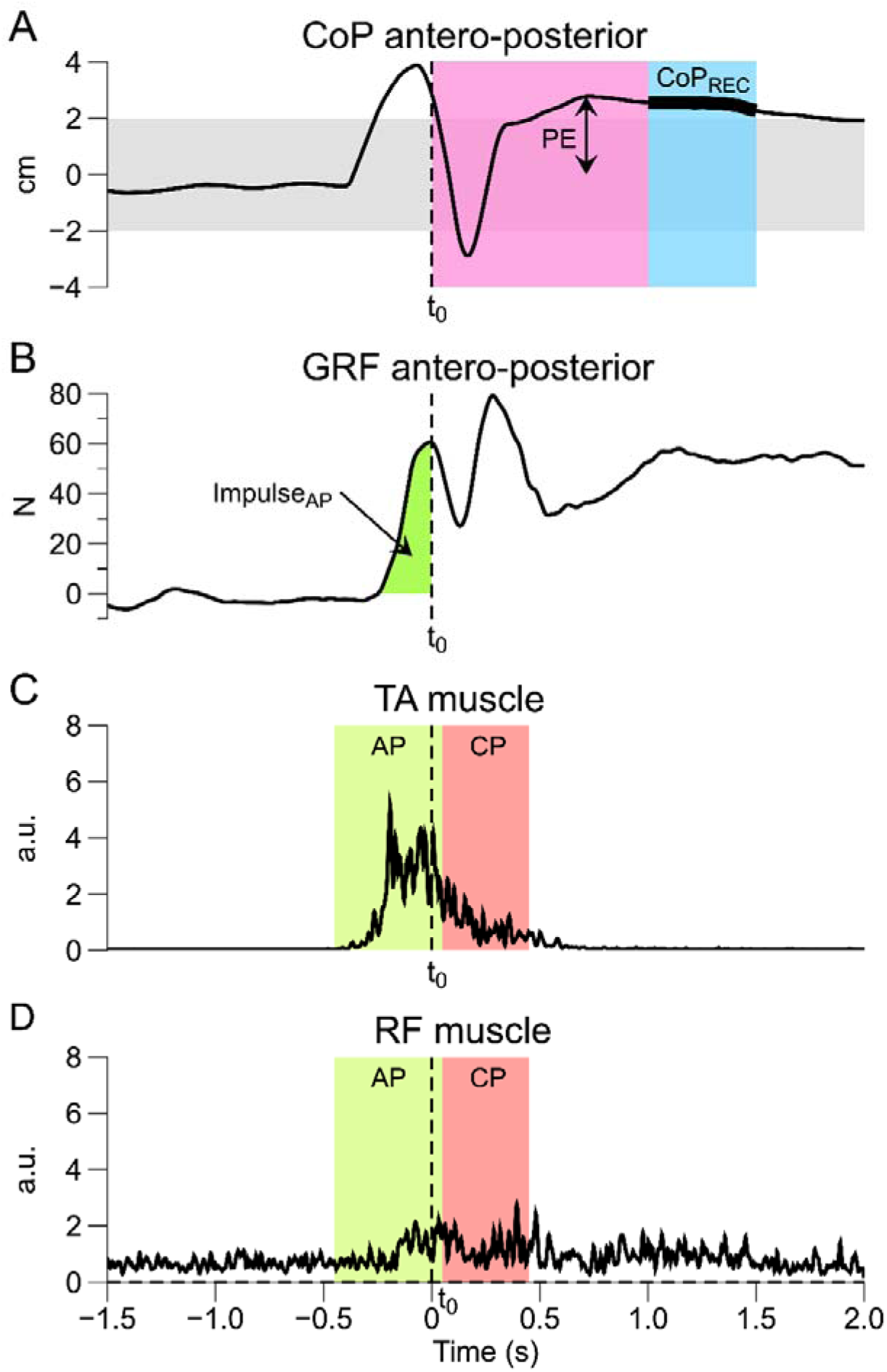
Aligned signals for a representative participant during a typical trial. The vertical dotted black line (t0) corresponds to the onset of the perturbation (i.e., button press). A: profile of the CoP in the antero-posterior direction, where the windows used to calculate CoP-related variables are highlighted. Performance error (PE) was computed as the peak backward (+) displacement of the CoP from the target position (horizontal black dotted line at y=0) during a 1s-window after t0 (PE window, magenta-shaded area). Recovery of CoP position (CoP_REC_; thick black line) was computed as the average position of the CoP during a 500-ms window (+1 to 1.5s after t0; light-blue shaded area). B: Ground reaction force signal in the antero-posterior direction. Highlighted in green is the Impulse_AP_, computed as the net (positive) integral of the signal before t0. C: Envelope (low-passed) EMG signal of the tibialis anterior (TA) during the postural task. Highlighted in light-green and light-red are the anticipatory (AP: t0-450:t0+50ms) and compensatory (CP: t0+50:t0+450) time-windows used for the analysis, respectively. D: Envelope (low-passed) EMG signal of the rectus femoris (RF) during the postural task. Highlighted in light-green and light-red are the anticipatory (AP: t0-450:t0+50ms) and compensatory (CP: t0+50:t0+450) time-windows used for the analysis, respectively.

Postural recovery following perturbation (CoP_REC_) was computed as the average position of the CoP during a 500-ms window (+1 to +1.5s after *t0*), relative to CoP_Target_. We used a signed CoP recovery metric because the direction of residual displacement is meaningful in this perturbation paradigm. The perturbation always acted in the backward direction, so positive CoP_REC_ values specifically reflect incomplete return toward the pre-perturbation posture rather than random oscillations. This measure reflects the effective recovery of the postural stability following the postural perturbation, not to be confused with the PE, which might be higher, in case of a compensatory forward movement following the absorption of the perturbation. To evaluate anticipatory movements by the CoP in preparation for the perturbation, we computed the integral (i.e., function *trapz* in MATLAB) over the positive signal of antero-posterior ground reaction force (Impulse_AP_) in the time window preceding *t_0_* (**Fig. 2B**).

#### EMG variables

Linear envelopes of pre-processed EMG signals were computed by applying a 50 Hz low-pass, 4^th^ order, zero-phase digital Butterworth filter. Pre-processed and envelope EMG signals for each muscle were inspected to discard unusable, poor-quality EMG recordings due to technical problems, abnormal signal-to-noise ratio or signal artifacts (i.e., electrode detachment or presence of sweat) or abrupt movements during the trials (e.g., stepping backward following the perturbation). EMG data were then normalized participant-wise by dividing each value by the mean late baseline (LB) activity for each muscle (Pienciak-Siewert et al., 2020), taken as the root-mean-square value of filtered EMG activity over a 900ms window around *t_0_* (*t_0_*-450:*t_0_*+450ms). Muscle activity was quantified as the root-mean-square value of normalized EMG over two separate fixed time-windows: anticipatory phase (AP: from -450 ms to + 50 ms respect to *t_0_*) and compensatory phase (CP; from +50 ms to + 450 ms respect to *t_0_*), as highlighted in **Fig. 2C** and **Fig. 2D**. Time-windows were chosen based on previous studies employing external perturbations (Santos et al., 2010; Piscitelli et al., 2017; Nardon et al., 2025) and on estimated onset in CoP movement during pilot tests for the current task.

Coactivation (CC Index) was quantified for each agonist-antagonist pair (e.g., TA vs. SOL) as the minimum value of normalized EMG (for each sampling point *i*, coactivation trace *CC Index_TA-SOL_* is computed from the individual muscle activity traces *TA* and *SOL* such that *CC Index_TA-SOL(_i)* = min[*TAi*, *SOLi*]). The resulting time-varying signal represents the magnitude of normalized EMG that is matched by the two opposing muscles (Thoroughman and Shadmehr, 1999; Gribble et al., 2003; Pienciak-Siewert et al., 2020). The representative coactivation value for each trial was calculated as the root-mean-square value of the time-varying coactivation signal during AP and CP time-windows.

### Statistical analysis

Jamovi statistical software (The jamovi project (2026). *jamovi* (Version 2.6.44) [Computer Software]. Retrieved from https://www.jamovi.org] was used for statistical analyses. All data in the text, Tables, and Figures – unless otherwise stated – are presented as mean ± 1 standard deviation (SD). For all comparisons, statistical significance was set at an α of 0.05. Data distribution (skewness and kurtosis) was assessed, the normality of the data was assessed by the Shapiro–Wilk test and confirmed by the inspection of density and Q–Q plots.

Fitting coefficients obtained from the PE adaptation curves (decay rate, *a*; asymptotic value, *b*; and initial value, *c*) were compared between groups using separate independent-samples t-tests for each coefficient and each experimental session (Day 1 and Day 2). Phase-averaged values of performance error (PE), *CoP_REC_*and *Impulse_AP_*, were analyzed with three-way repeated-measures ANOVAs, with *Phase* and *Session* (*Day 1*, *Day 2*) as within-subject factors and *Group* as a between-subjects factor. Primary outcomes were PE and CoP_REC_. Impulse_AP_, EMG and coactivation measures were considered exploratory. Considering the normalization method used and the possible variability in electrode placement across sessions, EMG and coactivation data were analyzed separately for each session (*Day 1*; *Day 2)* with a two-way repeated-measures ANOVAs, with *Phase* as within-subject factors and *Group* as a between-subjects factor. Assumptions of sphericity were explored using Mauchly’s test and controlled for using the Greenhouse–Geisser adjustment in instances where Mauchly’s test was significant (α <0.05). To test for adaptation, in the event of a significant interaction or main *Phase* effect, we made planned comparisons on the within-subject results for each group between the late baseline (LB), early adaptation (EA), late adaptation – start (LA_start_) and late adaptation – final block (LA_end_) phases. To compare the adaptation process between groups, we made planned comparisons between groups at LB, EA, LA_start_ and LA_end_ phases, for each day separately. Similarly, to test the effects of fatigue on the washout process, a 3-way repeated-measures ANOVA (*Phase* and *Session* as repeated measure factors) was used to assess differences in performance and postural variables across three experimental phases (LB, EW, LW). We used paired *t* tests for within-subject planned comparisons between phases and independent two-sample *t* tests for planned comparisons between groups. Statistic results within each ANOVA were then corrected for multiple comparisons using Bonferroni adjustment. We did not make across-ANOVA adjustments in p-values, due to the limited sample size and considering that Impulse_AP_, EMG and coactivation measures were considered as exploratory outcomes. When normality assumption was violated, ANOVAs were run on log-transformed data. Non-parametric independent samples t-test (Welch-test) was used for pairwise comparisons between groups if Levene’s test revealed unequal variance. Differing degrees of freedom in the ANOVA outputs are due to listwise deletion of participants at the variable level in case of a missing value.

## Results

Isometric fatiguing exercise duration on Day 1 was on average 4:45 ± 0:53 min:s (number of cycles: 7.1 ± 1.3), with a time reduction trend across bouts (bout1: 5:58 ± 1:47 min:s, bout2: 5:07 ± 1:52 min:s, bout3: 5:40 ± 2:51 min:s, bout4: 4:43 ± 2:45 min:s, bout5: 3:50 ± 1:39 min:s, bout6: 4:13 ± 2:55 min:s, bout7: 3:40 ± 1:49 min:s). Repeated measure ANOVA results indicate that MVC isometric force pre-exercise did not differ between groups on each session (Day1: FAT: 250.8 ± 83.7 N, NoFAT: 251.9 ± 91.6 N; Day2: FAT: 259.5 ± 78.8 N, NoFAT: 251.2 ± 97.6 N), and we did not observe significant changes in MVC values between sessions or *Session*\**Group* interaction (p > 0.05). The fatiguing exercise led to a significant drop in the average isometric force in ankle dorsiflexion for Fatigue group on Day 1, and no differences in both groups on Day 2 (Day1: FAT: -32 ± 4%, NoFAT: 1 ± 3%, p < 0.001; Day 2: FAT: -0.8 ± 2.2%, NoFAT: 0.4 ± 2.3%, p > 0.05)

### Postural performance

Participants in both groups reduced their performance error (PE) with practice, and initial PE values were lower on Day 2 than on Day 1 (**Fig. 3A**). The group-averaged fitting curves reported in **Fig. 3B** serve as a descriptive visualization of the average adaptation trajectories; statistical inference was based on fitting coefficients *a, b* and *c* – reported below with their 95% confidence intervals (CI—in square brackets) – which were estimated separately for each participant.

**Fig. 3.**
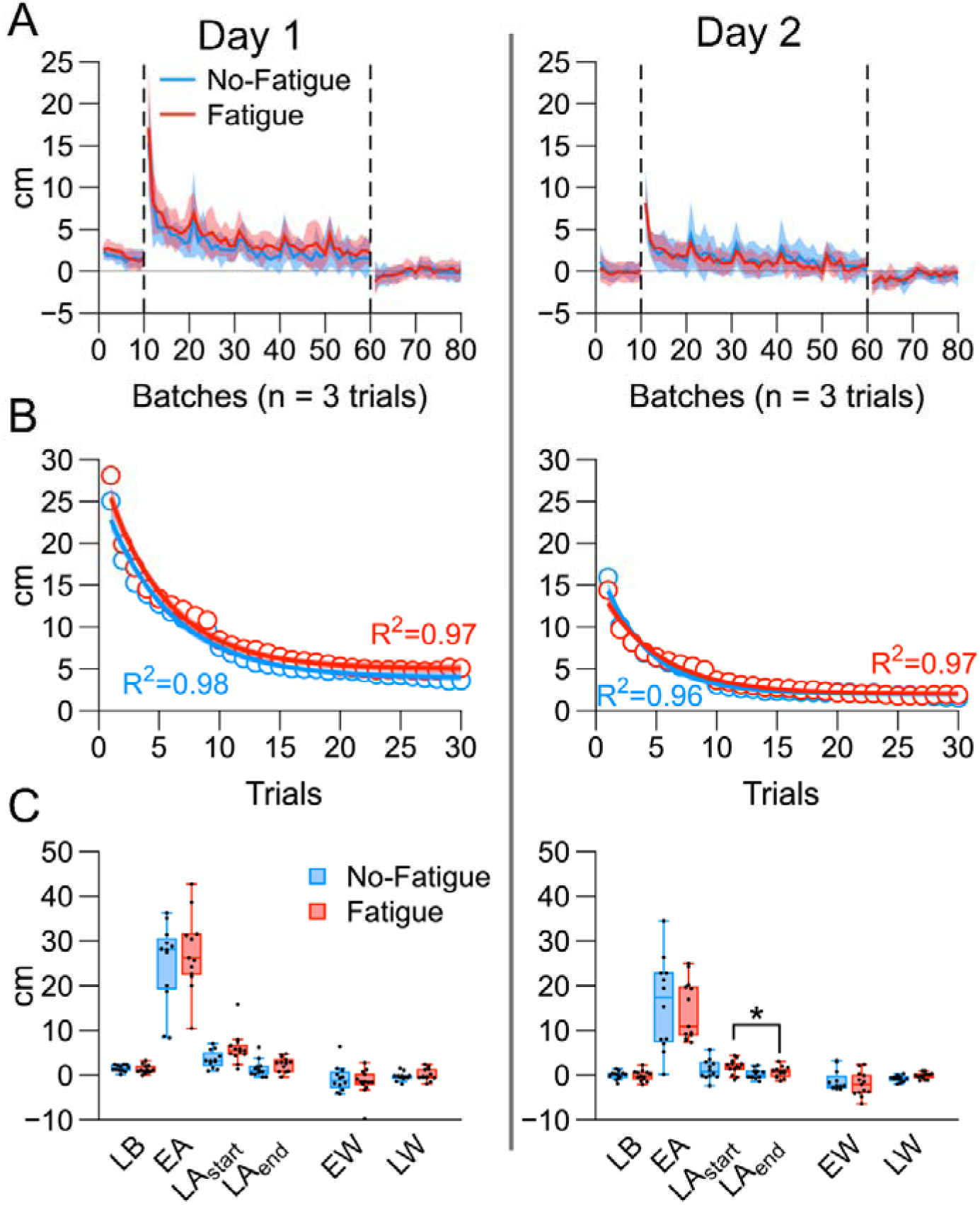
Group-averaged adaptation curves for performance error (PE), separated by day. A: PE throughout the experimental phases (baseline, adaptation, washout; separated by vertical dotted lines). Data are averaged in batches of n=3 trials and reported as mean (thick lines) ± 1 Standard Error (SE; shaded areas). Positive values represent a backward displacement. B: Visualization of average fitting curves (thick lines) and confidence intervals (CI) of the fittings (shaded areas) for PE group-averaged values (circles) during the first block of adaptation (30 trials). C: Box plots for PE across the considered experimental phases. Boxes depict data from 1^st^ to 3^rd^ quartile; whiskers represent minimum and maximum data points. Median is shown by the thick horizontal line and individual data points as black dots. * Significant effects (P < 0.05) are represented only in case of between-groups differences or within-group effects differing from the behavior of the other group. EA, early adaptation; EW, early washout; LAend, late adaptation final block; LAstart, late adaptation—start; LB, late baseline; LW, late washout.

On Day 1, the decay-rate coefficient *a* did not differ between groups (FAT: 0.43 [0.32, 0.53]; NoFAT: 0.35 [0.23, 0.47]; *t*_(23)_ = 1.05, *p* = 0.303, Cohen’s *d* = 0.42 [−0.38, 1.21]). The final steady-state coefficient *b* was higher in FAT than NoFAT (FAT: 5.23 [4.35, 6.11]; NoFAT: 3.53 [2.65, 4.41]; *t*_(23)_ = 2.98, *p* = 0.007, *d* = 1.19 [0.33, 2.04]). The initial-value coefficient *c* did not differ between groups (FAT: 28.46 [21.53, 35.40]; NoFAT: 35.71 [25.44, 45.98]; *t*_(23)_ = −1.30, *p* = 0.206, *d* = −0.52, 95% CI [−1.31, 0.28]).

On Day 2, coefficient *a* did not differ between groups (FAT: 0.37 [0.28, 0.46]; NoFAT: 0.62 [0.32, 0.93]; *t*_(23)_ = −1.83, *p* = 0.081, *d* = −0.73 [−1.54, 0.09]), nor did coefficient *b* (FAT: 2.02 [1.22, 2.81]; NoFAT: 1.92 [0.83, 3.01]; *t*(23) = 0.15, *p* = 0.879, *d* = 0.06, 95% CI [−0.72, 0.85]). For coefficient *c*, Levene’s test indicated unequal variances, so Welch’s test was used: the groups did not differ (FAT: 14.26 [10.28, 18.24]; NoFAT: 25.86 [14.33, 37.39]; Welch’s *t*_(13.65)_ = −2.09, *p* = 0.056, *d* = −0.85 [−1.68, 0.01]).

We observed a significant main effect for *Phase* (F_(1.13, 25.91)_ = 214.73, p < 0.001, η*_2p_* = 0.903) and *Session* (F_(1, 23)_ = 43.70, p < 0.001, η*_2p_* = 0.655) and a significant *Phase*Session* interaction (F_(1.16, 26.63)_ = 13.26, p < 0.001, η*_2p_* = 0.366). Planned comparisons in the adaptation phase confirmed that PE increased during EA and LA_start_ (p < 0.001) in both groups both on *Day 1* and *Day 2*, when compared with baseline (LB) values. Similarly, PE values during EA were significantly higher compared to all the successive adaptation time windows (LA_start_, LA_end_; p < 0.001) for both groups on both days. Finally, on Day 1 PE reduced significantly from LA_start_ to LA_end_ in both groups (p = 0.001; d = 0.998 [0.509, 1.47]). On Day 2, the decrease in PE values from LA_start_ to LA_end_ was significant only for the FAT group (p = 0.042; d = 0.796 [0.156, 1.41]; **Fig. 3C**). Main effect for *Group* was not significant (F_(1, 23)_ = 1.09, p = 0.308, η*_2p_* = 0.045).

Washout phase exhibited – as expected – an opposite trend, with a significant main effect for *Phase* (F_(1.15, 25.91)_ = 16.32, p < 0.001, η*^2^* = 0.415) and *Session* (F_(1, 23)_ = 10.87, p = 0.003, η*^2^* = 0.655). Pairwise comparisons revealed a significant increase in PE for both groups during EW (p = 0.006) and LW (p < 0.001), and a significant difference between EW and LW (p = 0.041) on Day 1. On Day 2, the increase in PE during EW was significant only for the FAT group (p = 0.033). Main effect for *Group* was not significant (F_(1, 23)_ = 0.14, p = 0.714, η*^2^* = 0.006).

#### Other postural variables

Additional variables were computed from CoP data, to capture the appreciable changes in the CoP profile throughout the adaptation process (**Fig. 4**). Postural recovery following perturbation (CoP_REC_) across the adaptation phase shows an appreciable reduction across time windows (**Fig. 5A**). The ANOVA confirmed a significant main effect for *Phase* (F_(3, 69)_ = 244.43, p < 0.001, η*_2p_* = 0.914), *Session* (F_(1, 23)_ = 58.00, p < 0.001, η*_2p_* = 0.716) and *Group* (F_(1, 23)_ = 12.90, p = 0.002, η*_2p_* = 0.359). Additionally, the ANOVA revealed a significant *Phase*Group* (F_(3, 69)_ = 2.79, p = 0.047, η*_2p_* = 0.108), *Session*Group* (F_(1, 23)_ = 4.85, p = 0.038, η*_2p_* = 0.174) and *Phase*Session* (F_(3, 69)_ = 12.41, p < 0.001, η*_2p_* = 0.351) interactions.

**Fig. 4.**
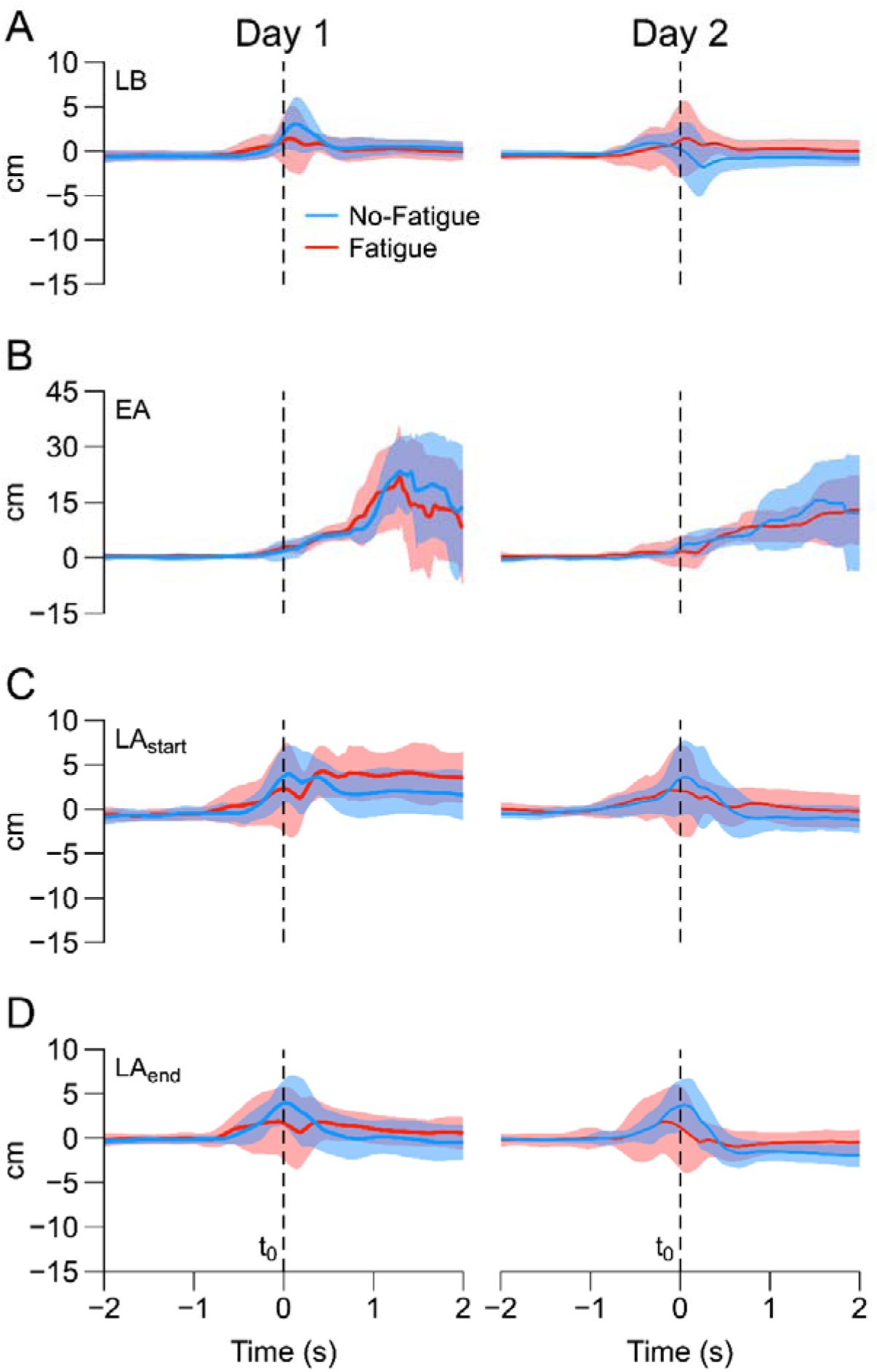
Group-averaged CoP_AP_ trajectory aligned at perturbation onset (t0) across the adaptation phase on Day 1 (left column) and Day 2 (right column). Thick lines represent means and shaded areas depict ± 1 Standard Deviation (SD). Vertical dashed lines are aligned at t0. Positive and negative values represent backward and forward displacement, respectively. A: Aligned CoP trajectories during late baseline (LB). B: Aligned CoP trajectories during early adaptation (EA). C: Aligned CoP trajectories during late adaptation – start (LA_start_). D: Aligned CoP trajectories during late adaptation – final block (LA_end_).

**Fig. 5.**
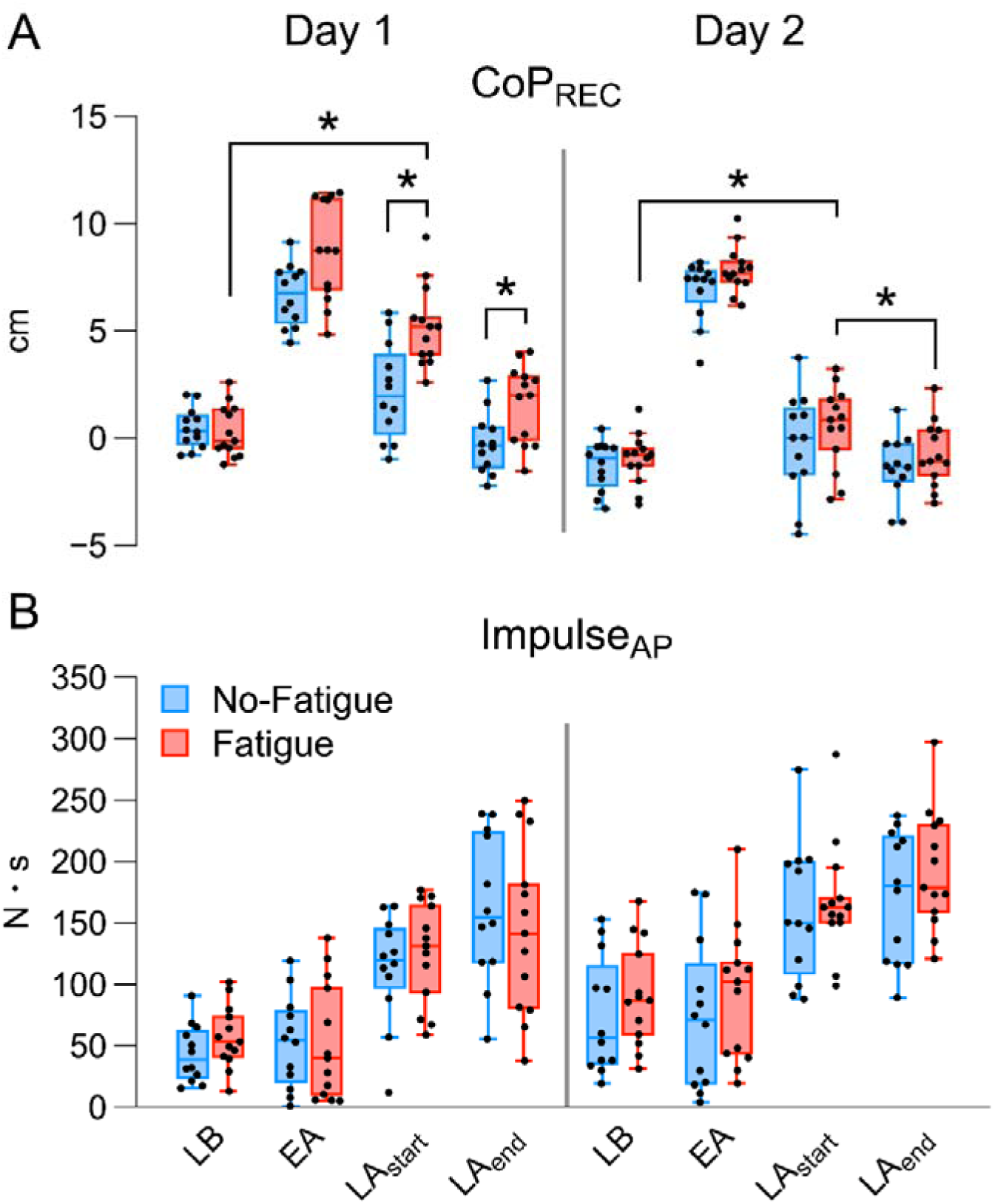
A: Box plots of CoP_REC_ across the adaptation phase. Values for Day 1 and Day 2 are positioned side-to-side, separated by a vertical line. B: Box plots of Impulse_AP_ across the adaptation phase. Values for Day 1 and Day 2 are positioned side-to-side, separated by a vertical line. Boxes depict data from 1^st^ to 3^rd^ quartile; whiskers represent minimum and maximum data points. Median is shown by the thick horizontal line and individual data points as black dots. * Significant between/within group effects (p < 0.05). EA: early adaptation, LA_end_: late adaptation – final block, LA_start_: late adaptation – start, LB: late baseline.

Planned comparisons in the adaptation phase revealed an increase of CoP_REC_ from LB to EA in both groups both on *Day 1* and *Day 2* (p < 0.001). The increase from LB to LA_start_ was significant only for FAT group, both on Day 1 (p < 0.001; d = 2.02 [1.037, 2.965]) and on Day 2 (p = 0.039; d = 0.809 [0.166, 1.428]). Differences between EA and LA_start_ and LA_end_ were significant for both groups on both days (p < 0.001). On Day 1, CoP_REC_ decreased in both groups from LA_start_ to LA_end_ (p < 0.001; d = -1.263 [-1.785, -0.727]), while on Day 2 the decrease was significant only in FAT group (p = 0.048; d = -0.763 [-1.373, -0.13]).

On Day 1, the planned comparisons between groups revealed higher values of CoP_REC_ for the FAT group at LA_start_ (p = 0.006; d = 1.459 [0.557, 2.336]) and LA_end_ (p = 0.04; d = 1.123 [0.264, 1.961]), while the comparison at EA did not reach significance threshold (p = 0.06; d = 1.064 [0.147, 1.945]).

On Day 2, none of the comparisons reached significance threshold (all p > 0.183; EA: d = 0.788 [-0.076, 1.623]; LA_start_: d = 0.354 [-0.451, 1.145]; LA_end_: d = 0.407 [-0.404, 1.2]).

During the washout phase we observed a significant main effect for *Phase* (F_(1.39, 29.24)_ = 68.57, p < 0.001, η*_2p_* = 0.766), *Session* (F_(1, 21)_ = 17.62, p < 0.001 η*^2^* = 0.456) and a *Phase*Session*Group* (F_(1.61, 33.75)_ = 4.365, p = 0.027, η*^2^* = 0.172) interaction. Pairwise comparisons on Day 1 revealed a significant reduction of CoP_REC_ during EW for both NoFAT (p < 0.001) and FAT group (p = 0.042). Values of CoP_REC_ during LB differed from LW only for NoFAT group (p < 0.001), and a significant difference between EW and LW (p < 0.001) was present only for NoFAT. On Day 2, a similar decrease in CoP_REC_ was observed at EW for both groups (p < 0.001) and a significant difference between EW and LW was present in both NoFAT and FAT group (p = 0.003 and p < 0.001, respectively). Comparisons between groups were not significant (see Supplementary Data).

Anticipatory movements by the body’s center of pressure in preparation for the perturbation, as estimated from Impulse_AP_ increased across the adaptation phase (**Fig. 5B**). The ANOVA confirmed a significant main effect for *Phase* (F_(2.37, 54.41)_ = 157.19, p < 0.001, η*^2^* = 0.872) and *Session* (F_(1, 23)_ = 33.22, p < 0.001, η*^2^* = 0.591). Planned comparisons in the adaptation phase revealed a similar increase of Impulse_AP_ in both groups from LB to LA_start_ and LA_end_ (both with p < 0.001), and from EA to LA_start_ and LA_end_ (both with p < 0.001) both on *Day 1* and *Day 2*. Similarly, we observed an increase of Impulse_AP_ from LA_start_ to LA_end_ on Day 1 (p = 0.029) and on Day 2 (p = 0.015). Planned comparisons between groups were not run in absence of a main *Group* effect or significant interactions.

During the washout phase we observed a significant main effect for *Phase* (F_(1.46, 33.62)_ = 28.204, p < 0.001, η*_2p_* = 0.551), *Session* (F_(1, 23)_ = 10.375, p = 0.004, η*_2p_* = 0.311) and a *Phase*Session* (F_(1.96, 45.1)_ = 30.49, p = < 0.001, η*_2p_* = 0.57) interaction. Pairwise comparisons revealed a significant increase in Impulse_AP_ for both groups during EW (p < 0.001) and LW (p < 0.001), and a significant difference between EW and LW (p = 0.031) on Day 1. On Day 2, pairwise comparisons did not reach statistical significance (see Supplementary Data). Planned comparisons between groups were not run in absence of a main *Group* effect or significant interactions.

### EMG activity

The number of participants included in the EMG and coactivation analyses varied by muscle, activation window, and session because invalid recordings were excluded listwise for the relevant analysis. The number of valid participants for each analysis ranged between 22 and 25. Exact numbers relative to each analysis are reported as Supplementary Data.

#### EMG activity of shank muscles

Across both days, tibialis anterior (TA) and soleus (SOL) exhibited clear modulation during anticipatory and compensatory phases (**Fig. 6**). TA activity increased from baseline to late adaptation, with the largest changes occurring during compensatory responses. Group differences emerged primarily at early adaptation (EA), where the FAT group showed higher TA activity (**Fig. 6A**).

**Fig. 6.**
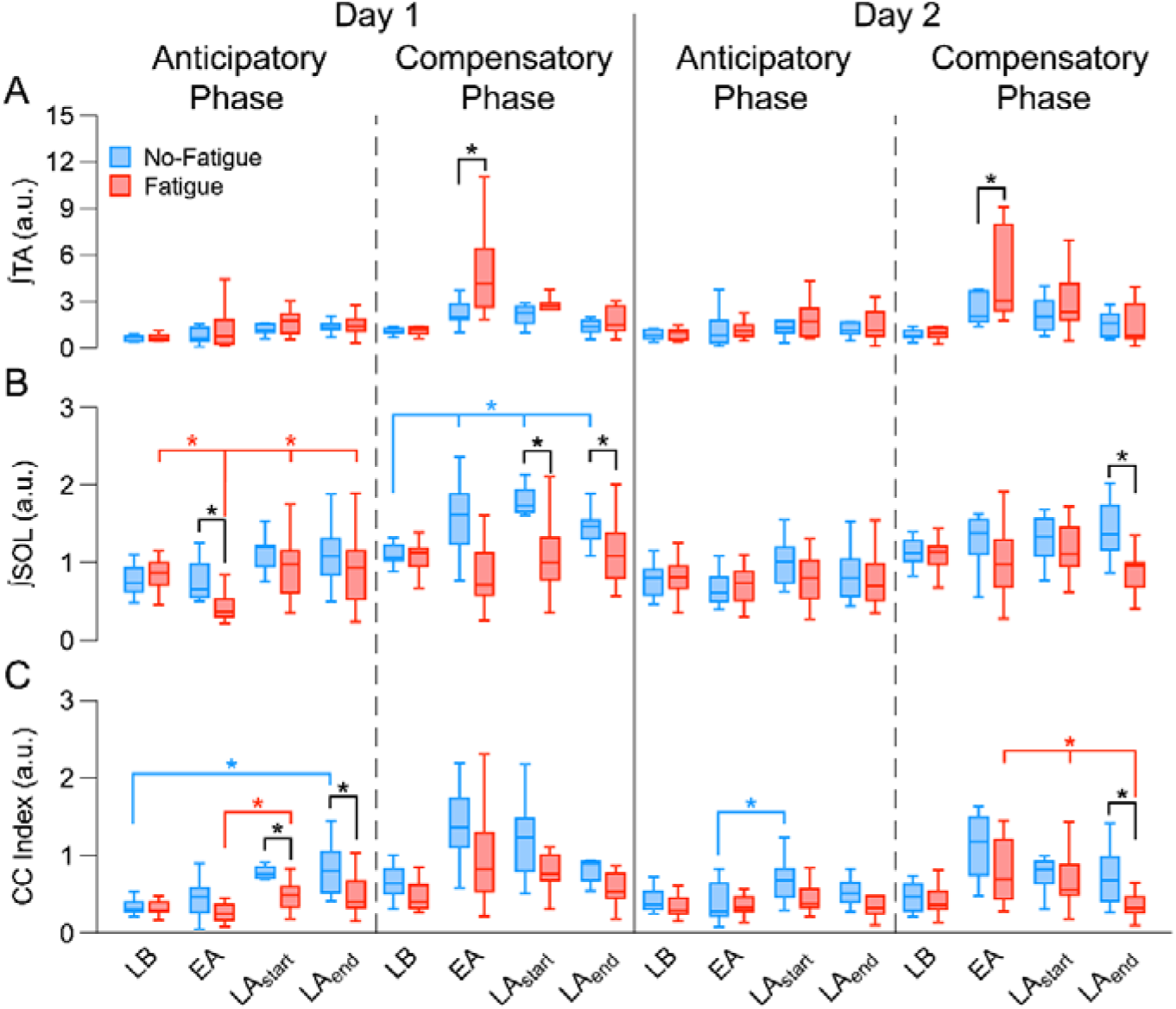
A: Box plots with EMG activity of tibialis anterior (TA) across the adaptation phase. B: Box plots with EMG activity of soleus (SOL) across the adaptation phase. C: Box plots with TA-SOL coactivation index (CC Index) across the adaptation phase. The thick vertical line separates results by session (Day 1, Day 2), while pre-determined time window (anticipatory (AP) and compensatory (CP) phases) are shown side by side for each day. Boxes depict data from 1^st^ to 3^rd^ quartile; whiskers represent minimum and maximum data points. Median is shown by the thick horizontal line. * Significant between/within group effects (p < 0.05). CC Index: coactivation index for tibialis anterior-soleus muscle pair, EA: early adaptation, LA_end_: late adaptation – final block, LA_start_: late adaptation – start, LB: late baseline, SOL: soleus, TA: tibialis anterior.

SOL displayed more pronounced between-group differences. On Day 1, the FAT group showed distinct anticipatory behavior, whereas compensatory activity differed mainly in the NoFAT group at late adaptation. On Day 2, SOL modulation was overall reduced, but a group difference persisted at the end of the adaptation phase (LA_end_), indicating that fatigue-induced coordination changes were retained during re-exposure (**Fig. 6B**).

#### TA-SOL muscle coactivation

Coactivation between TA and SOL increased from baseline to late adaptation on both days (**Fig. 6C**). On Day 1, the NoFAT group showed higher coactivation at late adaptation, suggesting a stabilization strategy relying on increased joint stiffness. On Day 2, coactivation followed a similar pattern, though between-group differences were smaller and limited to LA_end_. Detailed statistical analysis results for shank muscles EMG and CC Index are reported in **Table 2**.

**Table 2:**
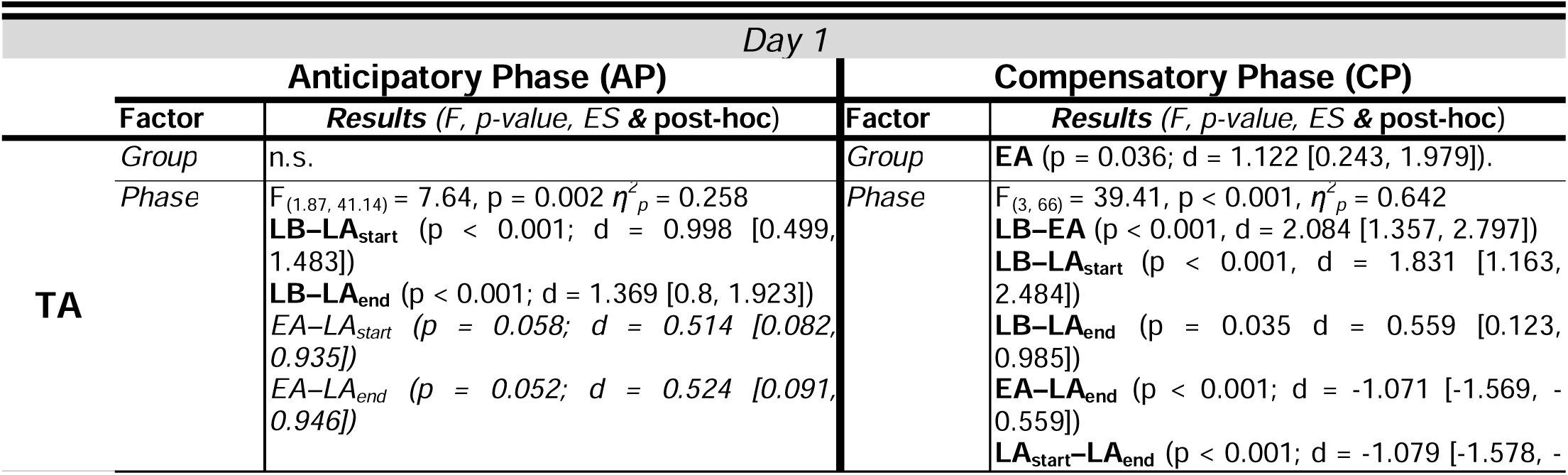
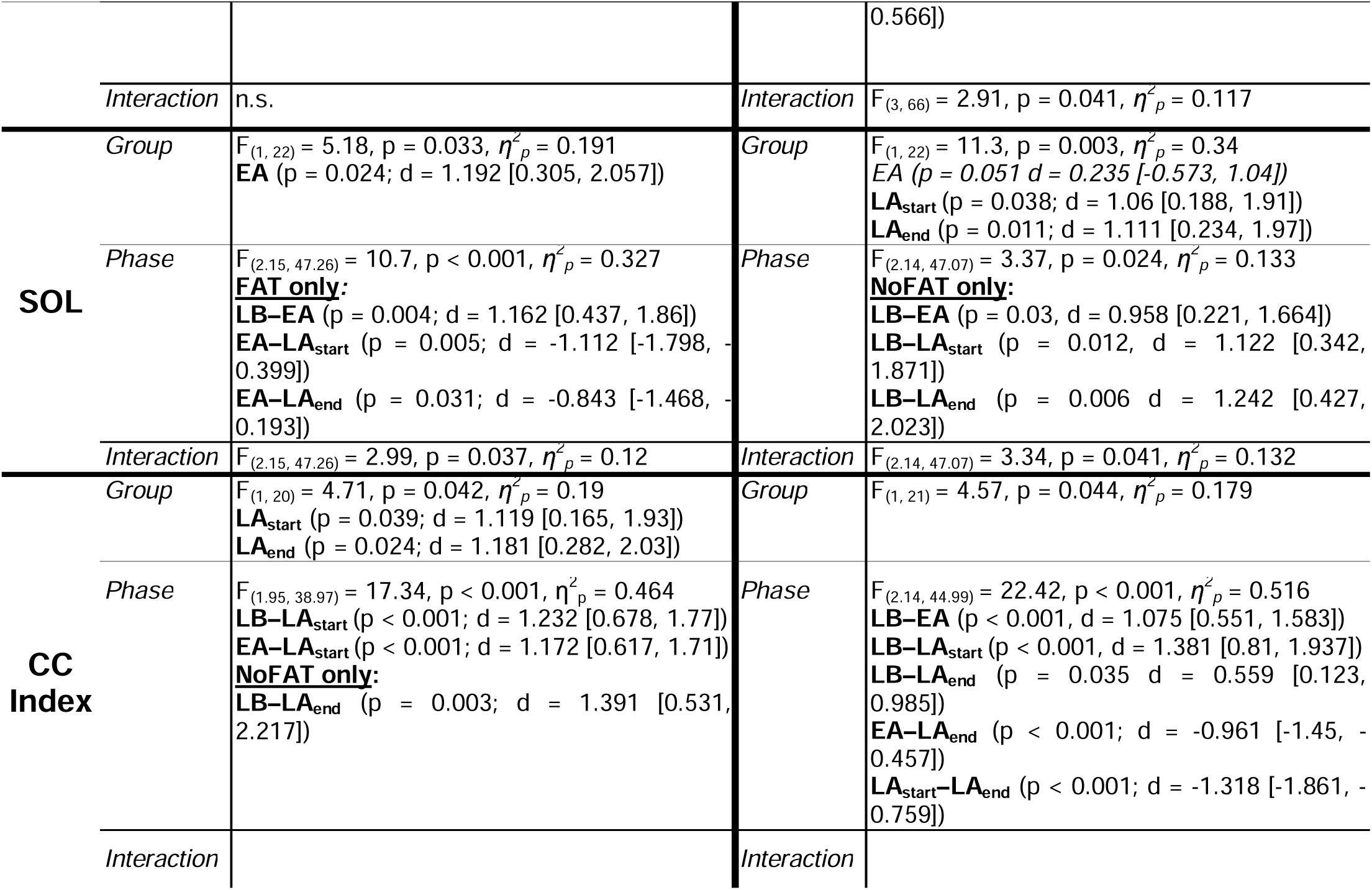
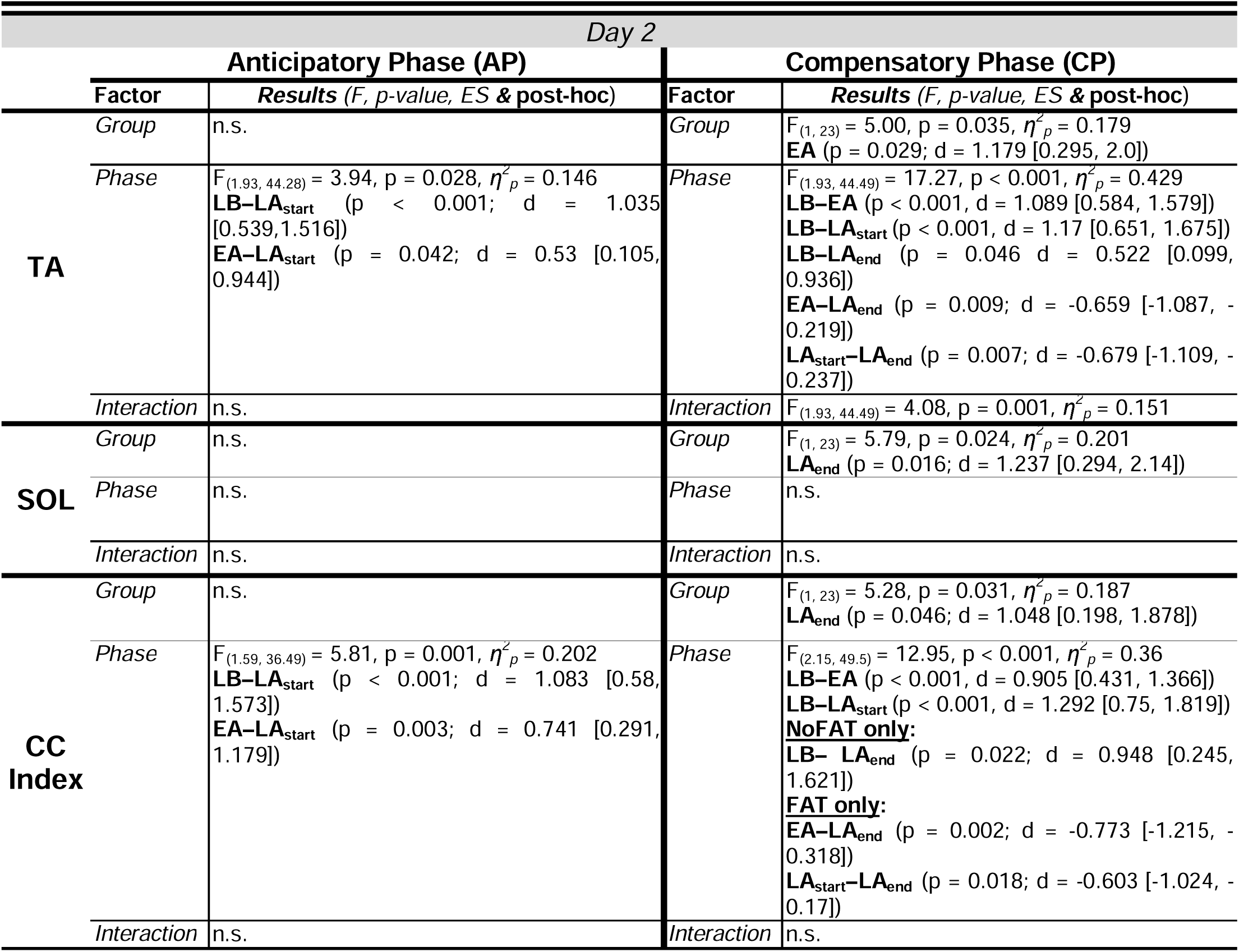
Statistical analysis results for shank muscles EMG – *tibialis anterior* (TA) and *soleus* (SOL) – and their coactivation index (CC Index) in the *adaptation phase*. Results for the *anticipatory phase* (AP) and *compensatory phase* (CP) are reported in side-by-side columns. Results are reported separately for each session (*Day 1, Day 2*). Abbreviations: ES: effect size, d: post-hoc effect size (Cohen’s *d*), η*^2^p*: ANOVA effect size (partial eta squared), LB: late baseline, EA: early adaptation, LA_start_: late adaptation – start, LA_end_: late adaptation – final block.

| *Day 1* |  |  |  |  |
|---|---|---|---|---|
|  | **Anticipatory Phase (AP)** |  | **Compensatory Phase (CP)** |  |
|  | **Factor** | ***Results*** *(F, p-value, ES* ***&*** **post-hoc**) | **Factor** | ***Results*** *(F, p-value, ES* ***&*** **post-hoc**) |
| **TA** | *Group* | n.s. | *Group* | **EA** (p = 0.036; d = 1.122 [0.243, 1.979]). |
|  | *Phase* | F<sub>(1.87, 41.14)</sub> = 7.64, p = 0.002 *η<sup>2</sup><sub>p</sub>* = 0.258<br>**LB–LA<sub>start</sub>** (p < 0.001; d = 0.998 [0.499, 1.483])<br>**LB–LA<sub>end</sub>** (p < 0.001; d = 1.369 [0.8, 1.923])<br>*EA–LA<sub>start</sub> (p = 0.058; d = 0.514 [0.082, 0.935])*<br>*EA–LA<sub>end</sub> (p = 0.052; d = 0.524 [0.091, 0.946])* | *Phase* | F<sub>(3, 66)</sub> = 39.41, p < 0.001, *η<sup>2</sup><sub>p</sub>* = 0.642<br>**LB–EA** (p < 0.001, d = 2.084 [1.357, 2.797])<br>**LB–LA<sub>start</sub>** (p < 0.001, d = 1.831 [1.163, 2.484])<br>**LB–LA<sub>end</sub>** (p = 0.035 d = 0.559 [0.123, 0.985])<br>**EA–LA<sub>end</sub>** (p < 0.001; d = -1.071 [-1.569, -0.559])<br>**LA<sub>start</sub>–LA<sub>end</sub>** (p < 0.001; d = -1.079 [-1.578, -0.566]) |
|  | *Interaction* | n.s. | *Interaction* | F<sub>(3, 66)</sub> = 2.91, p = 0.041, *η<sup>2</sup><sub>p</sub>* = 0.117 |
| **SOL** | *Group* | F<sub>(1, 22)</sub> = 5.18, p = 0.033, *η<sup>2</sup><sub>p</sub>* = 0.191<br>**EA** (p = 0.024; d = 1.192 [0.305, 2.057]) | *Group* | F<sub>(1, 22)</sub> = 11.3, p = 0.003, *η<sup>2</sup><sub>p</sub>* = 0.34<br>*EA (p = 0.051 d = 0.235 [-0.573, 1.04])*<br>**LA<sub>start</sub>** (p = 0.038; d = 1.06 [0.188, 1.91])<br>**LA<sub>end</sub>** (p = 0.011; d = 1.111 [0.234, 1.97]) |
|  | *Phase* | F<sub>(2.15, 47.26)</sub> = 10.7, p < 0.001, *η<sup>2</sup><sub>p</sub>* = 0.327<br>**FAT only:**<br>**LB–EA** (p = 0.004; d = 1.162 [0.437, 1.86])<br>**EA–LA<sub>start</sub>** (p = 0.005; d = -1.112 [-1.798, -0.399])<br>**EA–LA<sub>end</sub>** (p = 0.031; d = -0.843 [-1.468, -0.193]) | *Phase* | F<sub>(2.14, 47.07)</sub> = 3.37, p = 0.024, *η<sup>2</sup><sub>p</sub>* = 0.133<br>**NoFAT only:**<br>**LB–EA** (p = 0.03, d = 0.958 [0.221, 1.664])<br>**LB–LA<sub>start</sub>** (p = 0.012, d = 1.122 [0.342, 1.871])<br>**LB–LA<sub>end</sub>** (p = 0.006 d = 1.242 [0.427, 2.023]) |
|  | *Interaction* | F<sub>(2.15, 47.26)</sub> = 2.99, p = 0.037, *η<sup>2</sup><sub>p</sub>* = 0.12 | *Interaction* | F<sub>(2.14, 47.07)</sub> = 3.34, p = 0.041, *η<sup>2</sup><sub>p</sub>* = 0.132 |
| **CC Index** | *Group* | F<sub>(1, 20)</sub> = 4.71, p = 0.042, *η<sup>2</sup><sub>p</sub>* = 0.19<br>**LA<sub>start</sub>** (p = 0.039; d = 1.119 [0.165, 1.93])<br>**LA<sub>end</sub>** (p = 0.024; d = 1.181 [0.282, 2.03]) | *Group* | F<sub>(1, 21)</sub> = 4.57, p = 0.044, *η<sup>2</sup><sub>p</sub>* = 0.179 |
|  | *Phase* | F<sub>(1.95, 38.97)</sub> = 17.34, p < 0.001, η<sup>2</sup><sub>p</sub> = 0.464<br>**LB–LA<sub>start</sub>** (p < 0.001; d = 1.232 [0.678, 1.77])<br>**EA–LA<sub>start</sub>** (p < 0.001; d = 1.172 [0.617, 1.71])<br>**NoFAT only:**<br>**LB–LA<sub>end</sub>** (p = 0.003; d = 1.391 [0.531, 2.217]) | *Phase* | F<sub>(2.14, 44.99)</sub> = 22.42, p < 0.001, *η<sup>2</sup><sub>p</sub>* = 0.516<br>**LB–EA** (p < 0.001, d = 1.075 [0.551, 1.583])<br>**LB–LA<sub>start</sub>** (p < 0.001, d = 1.381 [0.81, 1.937])<br>**LB–LA<sub>end</sub>** (p = 0.035 d = 0.559 [0.123, 0.985])<br>**EA–LA<sub>end</sub>** (p < 0.001; d = -0.961 [-1.45, -0.457])<br>**LA<sub>start</sub>–LA<sub>end</sub>** (p < 0.001; d = -1.318 [-1.861, -0.759]) |
|  | *Interaction* |  | *Interaction* |  |
| *Day 2* |  |  |  |  |
|  | **Anticipatory Phase (AP)** |  | **Compensatory Phase (CP)** |  |
|  | **Factor** | ***Results*** *(F, p-value, ES* ***&*** **post-hoc**) | **Factor** | ***Results*** *(F, p-value, ES* ***&*** **post-hoc**) |
| **TA** | *Group* | n.s. | *Group* | F<sub>(1, 23)</sub> = 5.00, p = 0.035, *η<sup>2</sup><sub>p</sub>* = 0.179<br>**EA** (p = 0.029; d = 1.179 [0.295, 2.0]) |
|  | *Phase* | F<sub>(1.93, 44.28)</sub> = 3.94, p = 0.028, *η<sup>2</sup><sub>p</sub>* = 0.146<br>**LB–LA<sub>start</sub>** (p < 0.001; d = 1.035 [0.539,1.516])<br>**EA–LA<sub>start</sub>** (p = 0.042; d = 0.53 [0.105, 0.944]) | *Phase* | F<sub>(1.93, 44.49)</sub> = 17.27, p < 0.001, *η<sup>2</sup><sub>p</sub>* = 0.429<br>**LB–EA** (p < 0.001, d = 1.089 [0.584, 1.579])<br>**LB–LA<sub>start</sub>** (p < 0.001, d = 1.17 [0.651, 1.675])<br>**LB–LA<sub>end</sub>** (p = 0.046 d = 0.522 [0.099, 0.936])<br>**EA–LA<sub>end</sub>** (p = 0.009; d = -0.659 [-1.087, -0.219])<br>**LA<sub>start</sub>–LA<sub>end</sub>** (p = 0.007; d = -0.679 [-1.109, -0.237]) |
|  | *Interaction* | n.s. | *Interaction* | F<sub>(1.93, 44.49)</sub> = 4.08, p = 0.001, *η<sup>2</sup><sub>p</sub>* = 0.151 |
| **SOL** | *Group* | n.s. | *Group* | F<sub>(1, 23)</sub> = 5.79, p = 0.024, *η<sup>2</sup><sub>p</sub>* = 0.201<br>**LA<sub>end</sub>** (p = 0.016; d = 1.237 [0.294, 2.14]) |
|  | *Phase* | n.s. | *Phase* | n.s. |
|  | *Interaction* | n.s. | *Interaction* | n.s. |
| **CC Index** | *Group* | n.s. | *Group* | F<sub>(1, 23)</sub> = 5.28, p = 0.031, *η<sup>2</sup><sub>p</sub>* = 0.187<br>**LA<sub>end</sub>** (p = 0.046; d = 1.048 [0.198, 1.878]) |
|  | *Phase* | F<sub>(1.59, 36.49)</sub> = 5.81, p = 0.001, *η<sup>2</sup><sub>p</sub>* = 0.202<br>**LB–LA<sub>start</sub>** (p < 0.001; d = 1.083 [0.58, 1.573])<br>**EA–LA<sub>start</sub>** (p = 0.003; d = 0.741 [0.291, 1.179]) | *Phase* | F<sub>(2.15, 49.5)</sub> = 12.95, p < 0.001, *η<sup>2</sup><sub>p</sub>* = 0.36<br>**LB–EA** (p < 0.001, d = 0.905 [0.431, 1.366])<br>**LB–LA<sub>start</sub>** (p < 0.001, d = 1.292 [0.75, 1.819])<br>**NoFAT only:**<br>**LB– LA<sub>end</sub>** (p = 0.022; d = 0.948 [0.245, 1.621])<br>**FAT only:**<br>**EA–LA<sub>end</sub>** (p = 0.002; d = -0.773 [-1.215, -0.318])<br>**LA<sub>start</sub>–LA<sub>end</sub>** (p = 0.018; d = -0.603 [-1.024, -0.17]) |
|  | *Interaction* | n.s. | *Interaction* | n.s. |

#### EMG activity of thigh muscles

Rectus femoris (RF) showed consistent modulation across adaptation (**Fig. 7A**). Anticipatory RF activity increased from baseline to late adaptation, with the FAT group showing earlier changes (i.e., comparison: LB–LA_start_). During compensatory responses, both groups increased RF activity across adaptation, though reductions from early to late adaptation occurred differently within groups depending on the day.

**Fig. 7.**
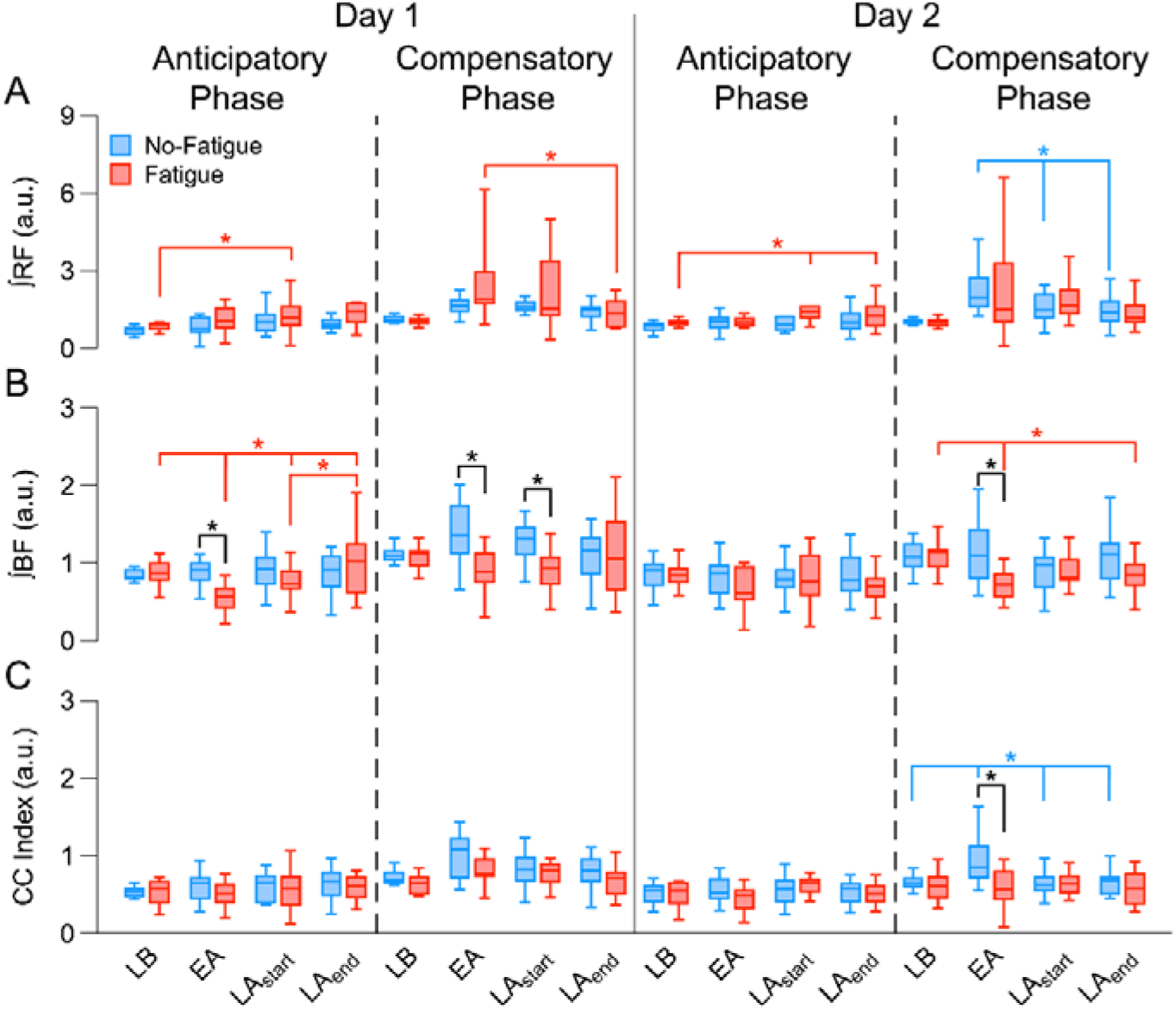
A: Box plots with EMG activity of rectus femoris (RF) across the adaptation phase. B: Box plots with EMG activity of biceps femoris (BF) across the adaptation phase. C: Box plots with RF-BF coactivation index (CC Index) across the adaptation phase. The thick vertical line separates results by session (Day 1, Day 2), while pre-determined time window (anticipatory (AP) and compensatory (CP) phases) are shown side by side for each day. Boxes depict data from 1^st^ to 3^rd^ quartile; whiskers represent minimum and maximum data points. Median is shown by the thick horizontal line. * Significant between/within group effects (p < 0.05). CC Index: coactivation index for rectus femoris-biceps femoris muscle pair, BF: biceps femoris, EA: early adaptation, LA_end_: late adaptation – final block, LA_start_: late adaptation – start, LB: late baseline, RF: rectus femoris.

Biceps femoris (BF) exhibited group-specific modulation (**Fig. 7B**). On Day 1, anticipatory BF activity modulated only in the FAT group, while compensatory activity showed clear between-group differences at EA and LA_start_. On Day 2, the modulation in BF was evident during the compensatory phase and only in the NoFAT group, which showed reductions from EA to LA_end_ and a persistent between-group difference at EA.

#### RF-BF muscle coactivation

Coactivation between RF and BF did not change during anticipatory phases. In the compensatory phase, however, coactivation increased from baseline to early and late adaptation on both days (**Fig. 7C**). Group differences appeared mainly at EA, and on Day 2 the NoFAT group showed progressive reductions from EA to late adaptation. Detailed statistical analysis results for thigh muscles EMG and CC Index are reported in **Table 3**.

**Table 3:**
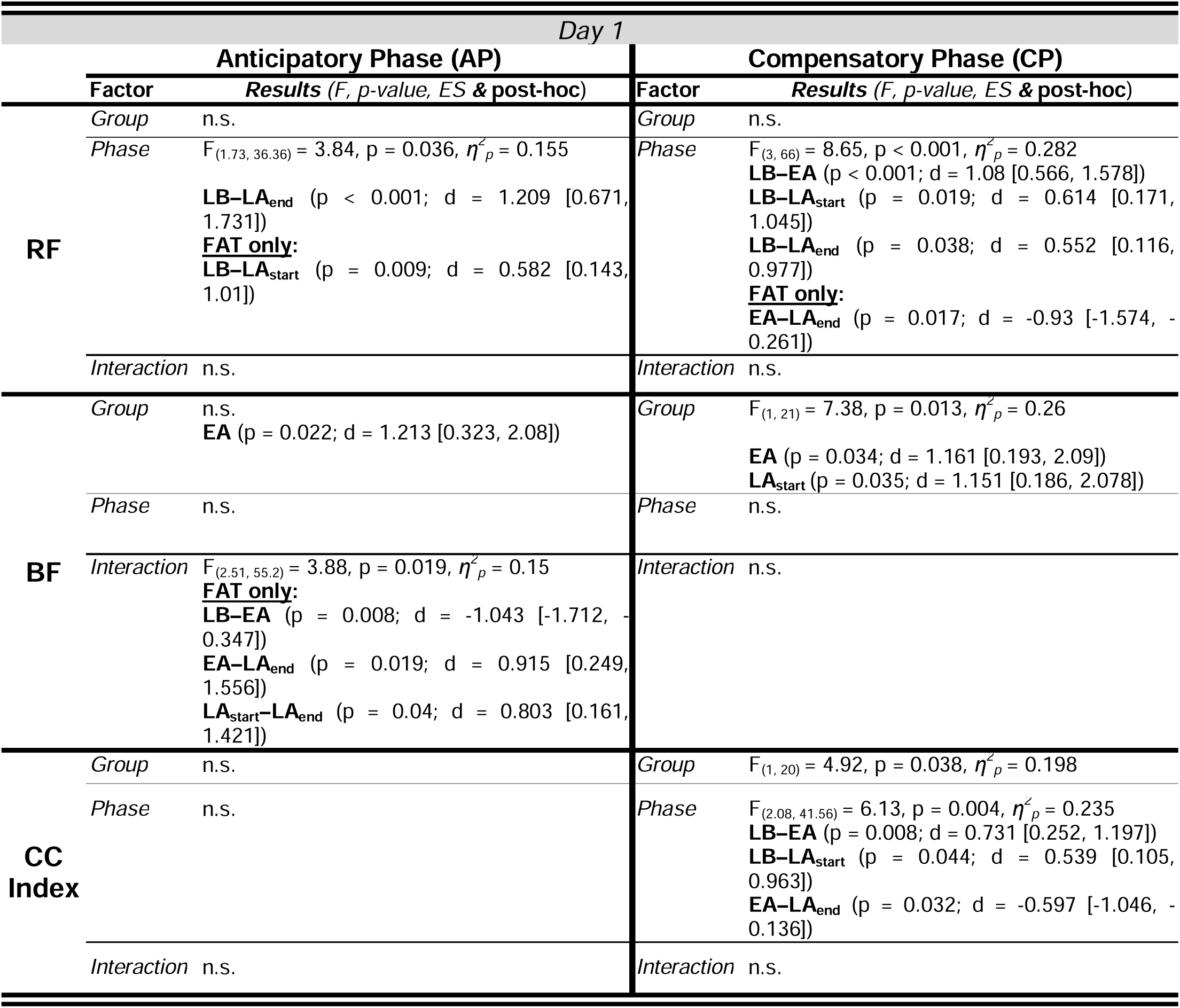
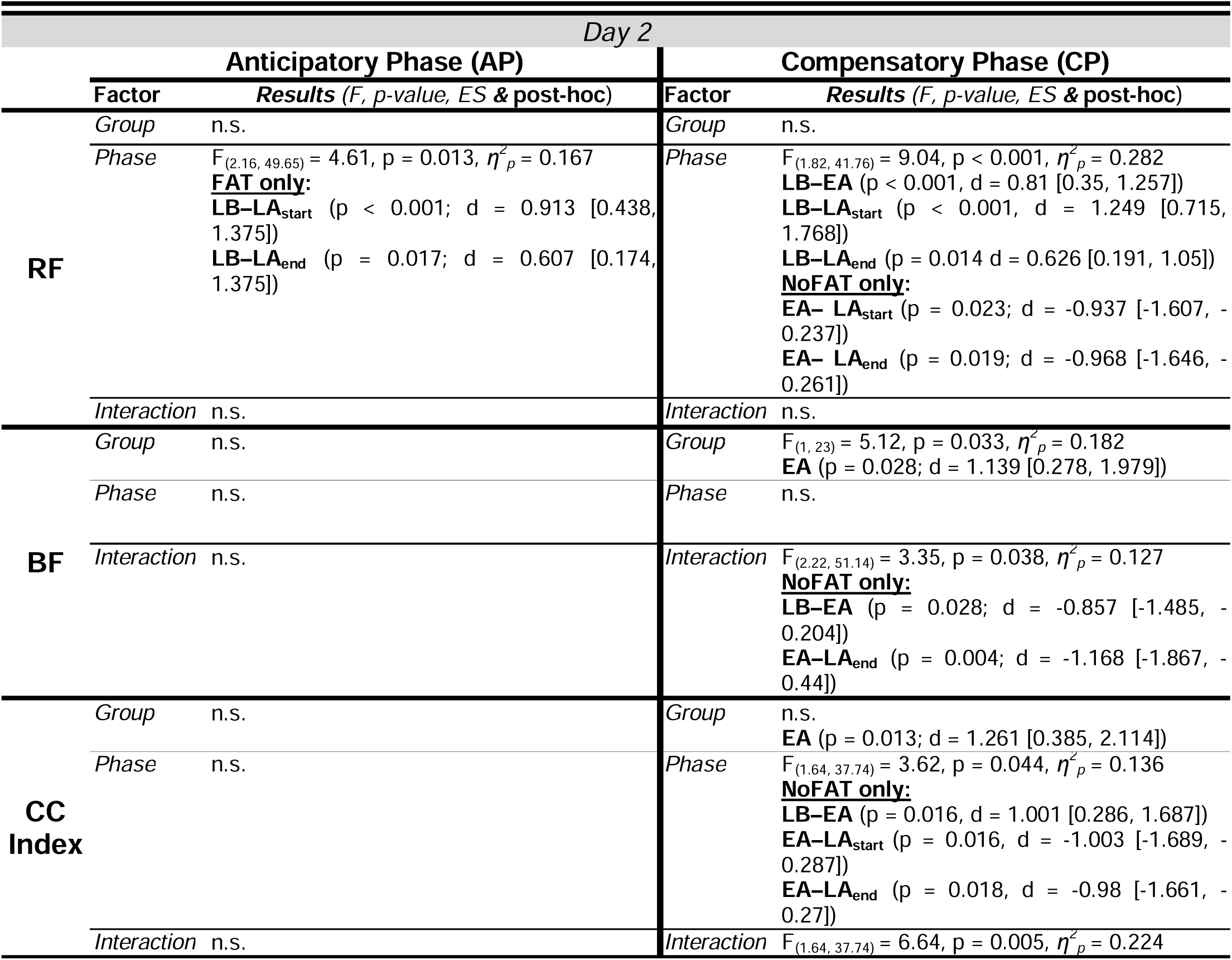
Statistical analysis results for thigh muscles EMG – *rectus femoris* (RF) and *biceps femoris* (BF) – and their coactivation index (CC Index) in the *adaptation phase.* Results for the *anticipatory phase* (AP) and *compensatory phase* (CP) are reported in side-by-side columns. Results are reported separately for each session (*Day 1, Day 2*). Abbreviations: ES: effect size, d: post-hoc effect size (Cohen’s *d*), η*^2^p*: ANOVA effect size (partial eta squared), LB: late baseline, EA: early adaptation, LA_start_: late adaptation – start, LA_end_: late adaptation – final block.

|  | Day 1 |  |  |  |
|---|---|---|---|---|
|  | **Anticipatory Phase (AP)** |  | **Compensatory Phase (CP)** |  |
|  | **Factor** | ***Results*** (*F, p-value, ES* ***&*** **post-hoc**) | **Factor** | ***Results*** (*F, p-value, ES* ***&*** **post-hoc**) |
| **RF** | *Group* | n.s. | *Group* | n.s. |
|  | *Phase* | F<sub>(1.73, 36.36)</sub> = 3.84, p = 0.036, *η*<sup>2</sup><sub>*p*</sub> = 0.155<br>**LB–LA<sub>end</sub>** (p < 0.001; d = 1.209 [0.671, 1.731])<br>**FAT only:**<br>**LB–LA<sub>start</sub>** (p = 0.009; d = 0.582 [0.143, 1.01]) | *Phase* | F<sub>(3, 66)</sub> = 8.65, p < 0.001, *η*<sup>2</sup><sub>*p*</sub> = 0.282<br>**LB–EA** (p < 0.001; d = 1.08 [0.566, 1.578])<br>**LB–LA<sub>start</sub>** (p = 0.019; d = 0.614 [0.171, 1.045])<br>**LB–LA<sub>end</sub>** (p = 0.038; d = 0.552 [0.116, 0.977])<br>**FAT only:**<br>**EA–LA<sub>end</sub>** (p = 0.017; d = -0.93 [-1.574, -0.261]) |
|  | *Interaction* | n.s. | *Interaction* | n.s. |
| **BF** | *Group* | n.s.<br>**EA** (p = 0.022; d = 1.213 [0.323, 2.08]) | *Group* | F<sub>(1, 21)</sub> = 7.38, p = 0.013, *η*<sup>2</sup><sub>*p*</sub> = 0.26<br>**EA** (p = 0.034; d = 1.161 [0.193, 2.09])<br>**LA<sub>start</sub>** (p = 0.035; d = 1.151 [0.186, 2.078]) |
|  | *Phase* | n.s. | *Phase* | n.s. |
|  | *Interaction* | F<sub>(2.51, 55.2)</sub> = 3.88, p = 0.019, *η*<sup>2</sup><sub>*p*</sub> = 0.15<br>**FAT only:**<br>**LB–EA** (p = 0.008; d = -1.043 [-1.712, -0.347])<br>**EA–LA<sub>end</sub>** (p = 0.019; d = 0.915 [0.249, 1.556])<br>**LA<sub>start</sub>–LA<sub>end</sub>** (p = 0.04; d = 0.803 [0.161, 1.421]) | *Interaction* | n.s. |
| **CC Index** | *Group* | n.s. | *Group* | F<sub>(1, 20)</sub> = 4.92, p = 0.038, *η*<sup>2</sup><sub>*p*</sub> = 0.198 |
|  | *Phase* | n.s. | *Phase* | F<sub>(2.08, 41.56)</sub> = 6.13, p = 0.004, *η*<sup>2</sup><sub>*p*</sub> = 0.235<br>**LB–EA** (p = 0.008; d = 0.731 [0.252, 1.197])<br>**LB–LA<sub>start</sub>** (p = 0.044; d = 0.539 [0.105, 0.963])<br>**EA–LA<sub>end</sub>** (p = 0.032; d = -0.597 [-1.046, -0.136]) |
|  | *Interaction* | n.s. | *Interaction* | n.s. |
|  | *Day 2* |  |  |  |
|  | **Anticipatory Phase (AP)** |  | **Compensatory Phase (CP)** |  |
|  | **Factor** | ***Results*** (*F, p-value, ES* ***&*** **post-hoc**) | **Factor** | ***Results*** (*F, p-value, ES* ***&*** **post-hoc**) |
| **RF** | *Group* | n.s. | *Group* | n.s. |
|  | *Phase* | F<sub>(2.16, 49.65)</sub> = 4.61, p = 0.013, *η*<sup>2</sup><sub>*p*</sub> = 0.167<br>**FAT only:**<br>**LB–LA<sub>start</sub>** (p < 0.001; d = 0.913 [0.438, 1.375])<br>**LB–LA<sub>end</sub>** (p = 0.017; d = 0.607 [0.174, 1.375]) | *Phase* | F<sub>(1.82, 41.76)</sub> = 9.04, p < 0.001, *η*<sup>2</sup><sub>*p*</sub> = 0.282<br>**LB–EA** (p < 0.001, d = 0.81 [0.35, 1.257])<br>**LB–LA<sub>start</sub>** (p < 0.001, d = 1.249 [0.715, 1.768])<br>**LB–LA<sub>end</sub>** (p = 0.014 d = 0.626 [0.191, 1.05])<br>**NoFAT only:**<br>**EA– LA<sub>start</sub>** (p = 0.023; d = -0.937 [-1.607, -0.237])<br>**EA– LA<sub>end</sub>** (p = 0.019; d = -0.968 [-1.646, -0.261]) |
|  | *Interaction* | n.s. | *Interaction* | n.s. |
| **BF** | *Group* | n.s. | *Group* | F<sub>(1, 23)</sub> = 5.12, p = 0.033, *η*<sup>2</sup><sub>*p*</sub> = 0.182<br>**EA** (p = 0.028; d = 1.139 [0.278, 1.979]) |
|  | *Phase* | n.s. | *Phase* | n.s. |
|  | *Interaction* | n.s. | *Interaction* | F<sub>(2.22, 51.14)</sub> = 3.35, p = 0.038, *η*<sup>2</sup><sub>*p*</sub> = 0.127<br>**NoFAT only:**<br>**LB–EA** (p = 0.028; d = -0.857 [-1.485, -0.204])<br>**EA–LA<sub>end</sub>** (p = 0.004; d = -1.168 [-1.867, -0.44]) |
| **CC Index** | *Group* | n.s. | *Group* | n.s.<br>**EA** (p = 0.013; d = 1.261 [0.385, 2.114]) |
|  | *Phase* | n.s. | *Phase* | F<sub>(1.64, 37.74)</sub> = 3.62, p = 0.044, *η*<sup>2</sup><sub>*p*</sub> = 0.136<br>**NoFAT only:**<br>**LB–EA** (p = 0.016, d = 1.001 [0.286, 1.687])<br>**EA–LA<sub>start</sub>** (p = 0.016, d = -1.003 [-1.689, -0.287])<br>**EA–LA<sub>end</sub>** (p = 0.018, d = -0.98 [-1.661, -0.27]) |
|  | *Interaction* | n.s. | *Interaction* | F<sub>(1.64, 37.74)</sub> = 6.64, p = 0.005, *η*<sup>2</sup><sub>*p*</sub> = 0.224 |

## DISCUSSION

The present study investigated how neuromuscular fatigue (NMF) of a postural muscle influences motor adaptation during a novel, whole-body postural control paradigm, and whether learning under fatigue produces persistent aftereffects during re-exposure to the perturbation (in the absence of neuromuscular fatigue).

Participants in both groups successfully adapted to the perturbation, reducing performance error across trials. This confirms that the fatigue-inducing protocol did not prevent behavioral adaptation, consistent with prior work showing that individuals can maintain task performance despite altered neuromuscular conditions (Nardon et al., 2024b). However, we observed that participants exposed to fatigue did not return to the initial posture following the postural perturbation throughout the adaptation phases. On the subsequent re-exposure to the task on a separate day – without being fatigued – participants in the FAT group showed similar aftereffects and a delayed return to baseline values. As expected, EMG patterns were altered by the presence of NMF, both in fatigued and non-fatigued muscles. Interestingly, these alterations persisted even in the subsequent re-exposure to the task, in absence of fatigue.

### Short-term effects on postural control

Contrary to our initial prediction, we did not observe between-group differences in task performance following localized NMF. Both groups showed an expected increase in error at the introduction of the perturbation followed by a progressive reduction across practice, consistent with classical adaptation processes driven by sensory prediction errors and iterative updating of internal models (Shadmehr et al., 2010; Krakauer et al., 2019). These findings indicate that participants exposed to NMF retained the capacity to adapt to the perturbation despite the altered neuromuscular state.

Nevertheless, a more detailed analysis of postural control variables revealed meaningful differences in the way participants exposed to fatigue managed the perturbation. Specifically, CoP_REC_ remained higher in the FAT group during the late stages of adaptation on Day 1, indicating a residual shift in the center-of-pressure following the perturbation. Importantly, this occurred despite comparable PE values between groups, suggesting that participants in the FAT group might have adopted alternative stabilization strategies that preserved task success while modifying whole-body postural behavior. These findings support the idea that motor adaptation under fatigue does not necessarily deteriorate, but rather relies on a reorganization of motor solutions to compensate for the altered neuromuscular state.

The absence of major deficits in PE may reflect the redundancy of the postural system. Unlike single-joint reaching paradigms where the fatigued muscle directly contributes to task execution, upright posture allows redistribution of motor commands across multiple joints and muscle groups (Bernstein, 1967; Latash, 2010; Singh and Latash, 2011). In the present study, although the tibialis anterior was selectively fatigued, participants could compensate through modified activation of proximal muscles and altered intermuscular coordination. Although in the present study we do not have directly assessed joint kinematics and kinetics, such compensatory behavior is consistent with previous evidence showing that the central nervous system exploits motor redundancy to preserve task-level goals under changing physiological constraints (Singh et al., 2010; Singh and Latash, 2011; Salomoni et al., 2019; Nardon et al., 2024b).

Interestingly, adaptation of anticipatory behavior appeared largely preserved under fatigue. Impulse_AP_ progressively increased across adaptation in both groups, indicating that participants learned to generate anticipatory shifts in the center of pressure before the perturbation. This pattern resembles the progressive emergence of anticipatory postural adjustments observed during repeated predictable perturbations (Santos et al., 2010; Piscitelli et al., 2017; Cesari et al., 2022). The preservation of this adaptation process suggests that NMF did not disrupt the ability to predict perturbation and generate feedforward responses (Kanekar et al., 2008). Rather, fatigue mainly affected the efficiency with which the perturbation was subsequently absorbed and stabilized. Alternatively, this measure could just represent a similar adaptation in the preparation for the voluntary button press, since it is not possible for this force measure to distinguish anticipation of the perturbation from the preparation of the voluntary action of button press itself.

These findings align with previous work showing dissociations between postural stability and task performance during fatigue. Nardon et al. (Nardon et al., 2024b) observed that participants exposed to lower-limb fatigue during an upper-limb force-field adaptation task maintained overall postural stability despite larger reaching errors, suggesting prioritization of balance-related control. In the present study, where postural control itself represented the task outcome, participants similarly maintained overall performance but displayed altered recovery dynamics, indicating that stabilization was achieved through different motor solutions. Together, these findings support the notion that the CNS flexibly reallocates control strategies under fatigue to preserve successful interaction with the environment.

Although CoP_REC_ captures direction-specific recovery dynamics, it does not distinguish residual displacement from potential overshoot. Another possible explanation for the altered recovery behavior observed in the FAT group relates to changes in proprioceptive acuity induced by fatigue. Localized fatigue of ankle musculature has been shown to impair force sense and joint position sense (Forestier et al., 2002; Vuillerme and Boisgontier, 2008), potentially reducing the accuracy of sensory feedback available for online corrections. Since postural stabilization following perturbation relies heavily on integration of proprioceptive information from the ankle joint (Peterka, 2002; Mohapatra et al., 2012), degraded sensory feedback may have contributed to the persistent backward displacement observed after perturbation release. Participants may therefore have relied more strongly on conservative stabilization strategies aimed at minimizing instability rather than limiting residual CoP displacements.

### Long-term effects on postural control

One of the aims of this study was to determine whether adaptation under fatigue produces persistent aftereffects during re-exposure to the perturbation. Although between-group differences in PE were again absent on Day 2, and the between-group estimates for CoP_REC_ patterns do not establish a significant difference, within-group patterns of CoP_REC_ remained elevated during adaptation compared with baseline, possibly suggesting that the FAT group continued to exhibit altered postural recovery dynamics. We acknowledge that this comparison alone cannot show that a fatigue-related group difference persisted even after recovery from the physiological effects of fatigue itself.

Nonetheless, these findings can be considered alongside prior work suggesting that fatigue may influence motor adaptation and subsequent learning (Takahashi et al., 2006; Huang et al., 2011; Branscheidt et al., 2019). Specifically, Takahashi et al. (Takahashi et al., 2006) demonstrated that adaptation performed under muscle fatigue modified aftereffects during subsequent testing, suggesting that the nervous system incorporated the fatigued state into the learned motor representation. Similarly, Branscheidt et al. (Branscheidt et al., 2019) demonstrated that learning under fatigue can produce persistent detrimental effects on subsequent motor-skill acquisition, even after recovery from the acute physiological effects of fatigue. Our findings extend this concept to a more ecological whole-body postural task, suggesting that fatigue-related motor solutions can possibly persist even in highly redundant systems.

Importantly, the persistence of within-group altered CoP_REC_ on Day 2 for FAT group despite recovery from acute fatigue may suggest that participants did not simply revert to their original strategy on Day 2. Instead, they appeared to recall and reuse motor solutions established during Day 1 in a fatigued condition. From a motor learning perspective, this may indicate that the CNS encoded not only the external perturbation but also the sensorimotor context in which adaptation occurred. State-dependent learning mechanisms have previously been proposed in locomotor and reaching paradigms, where internal models are partially linked to the physiological state experienced during practice (Bock et al., 2001; Torres-Oviedo and Bastian, 2012).

Interestingly, the persistence of altered stabilization strategies occurred despite largely similar anticipatory behavior between groups on Day 2. Impulse_AP_ adaptation followed comparable trajectories in both groups, indicating that predictive aspects of the task were retained similarly. The divergence may therefore indicate that the observed group patterns were more apparent in post-perturbation recovery than in anticipatory behavior, but the nonsignificant group comparisons and limited precision warrant caution. The EMG findings may provide complementary evidence about session-specific muscle-activation and coactivation patterns modulation during re-exposure.

### Fatigue-Induced effects on EMG Activation and Coactivation Patterns

The EMG results support the idea that adaptation under fatigue relied on substantial reorganization of neuromuscular control. The most evident effects involved both the fatigued tibialis anterior muscle and muscles not directly exposed to the fatiguing protocol, indicating that fatigue-related adaptations extended beyond the local muscular level.

During Day 1, the FAT group exhibited altered activation patterns in both anticipatory and compensatory phases, particularly involving TA, soleus, and biceps femoris muscles. In the compensatory phase, TA activity was significantly greater in the FAT group during early adaptation on both days, suggesting an increased neural drive to the fatigued muscle to compensate for reduced force-generating capacity. Similar increases in EMG amplitude following fatigue have been widely reported and are generally interpreted as evidence of compensatory recruitment strategies aimed at preserving force output despite reduced contractile efficiency (Taylor et al., 2000; Enoka and Duchateau, 2008; Enoka et al., 2011; Carroll et al., 2017).

At the same time, soleus activity and TA-SOL coactivation patterns differed between groups, particularly during Day 1. Participants in the NoFAT group progressively increased coactivation across adaptation, whereas the FAT group showed reduced coactivation levels during late adaptation. Coactivation is often indirectly interpreted as a strategy to increase joint stiffness and reduce movement variability when adapting to unstable or uncertain conditions (Thoroughman and Shadmehr, 1999; Osu et al., 2002; Gribble et al., 2003; Pienciak-Siewert et al., 2020; Martino et al., 2023). Reduced coactivation in the FAT group may therefore reflect an inability or unwillingness to further increase ankle stiffness due to the altered physiological state of the dorsiflexor muscles. Similar fatigue-induced reduction in muscle coactivation has been already documented (Nardon et al., 2025).

Furthermore, results are in line with findings from a recent study (Nardon et al., 2024b), where participants exposed to fatigue during motor adaptation in a standing reaching task with the arm showed decreased agonist-antagonist muscle co-contraction.

Instead, participants exposed to fatigue appeared to redistribute stabilization demands toward proximal musculature. This interpretation is indirectly supported by the modulation observed in rectus femoris and biceps femoris activity. Although we did not record trunk or hip kinematics in this study, modulations in muscle activity and coactivation might be explained by a proximal redistribution of muscle activity, similar to the “hip strategy” commonly described during challenging balance conditions or reduced ankle reliability (Horak and Nashner, 1986; Winter et al., 1998). Since ankle proprioception and force production were likely compromised by fatigue, greater reliance on proximal muscles may have represented a compensatory mechanism to maintain overall task success.

Importantly, several of these session-specific differences in EMG modulation persisted during Day 2 despite the absence of acute fatigue. The persistence of altered TA activity, BF activation, and coactivation patterns suggests that participants retained aspects of the neuromuscular strategy acquired during the fatigued condition. This finding reinforces the idea that adaptation under fatigue involves learning of a modified motor solution rather than only transient physiological compensation.

The persistence of altered coactivation patterns is particularly interesting from a motor control perspective. Previous work has shown that coactivation can decrease as internal models become more accurate and movement uncertainty declines (Thoroughman and Shadmehr, 1999; Gribble et al., 2003), while other studies suggest that muscle coactivation might be linked to inherent characteristics of the task (Darainy and Ostry, 2008; Martino et al., 2023). In the present study, however, the FAT group maintained different coactivation behavior respect to the NoFAT group, even after re-exposure, suggesting that the nervous system may have continued to treat the task as requiring a more conservative stabilization strategy. Such persistence may reflect altered estimates of sensorimotor reliability resulting from the initial fatigued experience. Interestingly, similar aftereffects on consolidated motor strategies during re-exposure to the task have been recently observed in participants who initially learned in presence of pain (Salomoni et al., 2019). Because EMG amplitudes were normalized within-session to the late-baseline activity in the present study, persistent differences across days should therefore be cautiously interpreted as differences in relative EMG modulation, not as evidence of unchanged absolute activation strategies.

Together, the EMG findings indicate that NMF influenced both local muscle activation and whole-body coordination strategies. Rather than preventing adaptation, fatigue altered the neuromuscular pathways through which adaptation was achieved. These results highlight the flexibility of the motor system in reorganizing control strategies under altered physiological constraints while preserving behavioral performance.

### Methodological considerations

Some limitations should be considered when interpreting the present findings. First, only a subset of postural muscles was monitored through surface EMG. Inclusion of trunk and hip musculature could provide a more comprehensive understanding of compensatory strategies during adaptation under fatigue. Furthermore, anticipatory movements by the CoP (Impulse_AP_) and EMG activity are consistent with postural adjustments occurring in preparation for the perturbation. However, because participants triggered the perturbation with a voluntary button press, we cannot exclude that these measures may also represent activity associated with preparing for or performing that action, rather than solely reflect predictive postural control. It is also worthwhile to note that our interpretations about redistribution of stabilization demands toward proximal musculature are limited to EMG recordings and should be confirmed by future studies accompanying muscle activity recordings with joint kinematics, joint moments, or direct estimates of joint stiffness.

Second, while the present task was intentionally designed to increase ecological validity compared with single-joint paradigms, its laboratory-based nature still differs from real-world balance tasks involving locomotion or multidirectional perturbations. Future investigation should determine whether the observed fatigue-induced alterations occur during ecological, more complex athletic movements and how differing fatiguing modalities could modify their interpretation.

Third, the relatively small sample and unequal sex distribution between groups should be considered when interpreting the findings. Sex distribution was not stratified; however, literature suggests minimal sex differences in dorsiflexor fatigability. Although the primary results were similar in an exploratory sensitivity analysis including sex, the sample size limits the precision of this assessment. Future studies with larger, more balanced samples are needed to confirm the findings and examine potential sex-related differences. We also acknowledge that, although perturbation magnitude was normalized to participants’ body mass, this procedure does not necessarily eliminate all potential effects of body dimensions on postural biomechanics. In addition, although sessions were separated by 48–72 hours to minimize residual fatigue and baseline MVC in both groups were not significantly different on Day 2, we cannot completely exclude subtle carry-over effects related to consolidation or contextual learning. Future studies should investigate whether the persistence of fatigue-related motor solutions extends over longer timescales and whether similar effects emerge in older adults or clinical populations with impaired postural control.

### Conclusions

Participants exposed to localized neuromuscular fatigue of the ankle dorsiflexors preserved behavioral adaptation to a novel postural perturbation but adopted distinct neuromuscular coordination strategies during practice, characterized by altered activation and coactivation patterns involving both fatigued and non-fatigued muscles. Several of these differences persisted during re-exposure 48–72 hours later, suggesting next-session persistence of altered stabilization strategies even in the absence of acute fatigue. Future work should examine whether such strategy-level changes generalize across different tasks and contexts, how they interact with sensory feedback alterations, and whether they persist over longer timescales.

These findings may provide a basis for further investigating the role of fatigue during motor learning in rehabilitation and athletic training contexts. In real-world settings, motor learning often occurs under varying levels of fatigue, and the conditions under which practice takes place may influence not only immediate performance but also the persistence of adapted movement strategies. However, whether the observed changes extend to more complex athletic movements, prolonged exercise, or different types of fatigue remains to be established. Future studies should therefore investigate whether fatigue-related adaptations are retained and transferred to functional tasks, and whether they have beneficial or detrimental consequences for movement control under fatigued and non-fatigued conditions.

## Appendix A. Supplementary Data

Supplementary Material contains additional figures and tables (**S1–S6**) presenting detailed results and statistical outputs for variables during the washout phase:

**Figure S1**: Results for *CoP_REC_ and Impulse_AP_*across the washout phase.

**Figure S2**: Results for EMGs of shank muscles (TA, SOL, *CC Index_TA-SOL_*) across the washout phase.

**Table S3**: ANOVA results for shank muscles EMG – *tibialis anterior* (TA) and *soleus* (SOL) – and their coactivation index (CC Index) in the washout phase.

**Figure S4**: Results for EMGs of thigh muscles (RA, BF, *CC Index_RF-BF_*) across the washout phase.

**Table S5**: ANOVA results for thigh muscles EMG – *rectus femoris* (RF) and *biceps femoris* (BF) – and their coactivation index (CC Index) in the washout phase.

**Table S6**: Number of participants included in each repeated measures ANOVA analysis for EMG measures after listwise exclusion.

The same material can be found on OSF: https://doi.org/10.17605/OSF.IO/A2XV4

## CRediT authorship contribution statement

**Mauro Nardon:** Writing – original draft, Investigation, Software, Data curation, Formal analysis, Conceptualization, Visualization. **Cristiano Alessandro:** Writing – review & editing, Conceptualization. **Tarkeshwar Singh:** Writing – review & editing, Formal analysis, Supervision, Conceptualization. **Matteo Bertucco:** Writing – review & editing, Investigation, Formal analysis, Visualization, Supervision, Conceptualization, Project administration.

## Acknowledgments

We would like to thank Dr. Francesco Pascucci for providing technical assistance in the implementation of the Arduino board connection and its integration with the experimental setup.

## Declaration of generative AI and AI-assisted technologies

During the preparation of this work, the authors used ChatGPT (OpenAI) to assist with the graphical representation of the experimental setup. The authors verified and edited the resulting image to ensure that it accurately represents the experimental apparatus and procedures and take full responsibility for the content of the published article.

## Data availability statement

De-identified original datasets from this study are publicly available on the OSF repository (**10.17605/OSF.IO/A2XV4**).

## Author contributions

M.N., M.B., and T.S. conceived and designed research; M.N. and M.B. performed experiments; M.N., M.B., and T.S. analyzed data; M.N., C.A., M.B., and T.S. interpreted results of experiments; M.N. and M.B. prepared figures; M.N., M.B., and T.S. drafted manuscript; M.N., C.A., M.B., and T.S. edited and revised manuscript; M.N., C.A., M.B., and T.S. approved final version of manuscript.

## Funding

This research did not receive any specific grant from funding agencies in the public, commercial, or not-for-profit sectors.

## Declaration of competing interest

The authors declare that they have no known competing financial interests or personal relationships that could have appeared to influence the work reported in this paper.

## Abbreviations

AP: Anticipatory phase
APAs: Anticipatory postural adjustments
BF: Biceps femoris
CC Index: Coactivation index
CNS: Central nervous system
CoP_AP_: CoP position in the antero-posterior direction
CoP_Feedback_: Online visual feedback of CoP
CoP_REC_: Postural recovery following the perturbation
CoP_Target_: CoP target position
CP: Compensatory phase
CPAs: compensatory postural adjustments
EA: Early adaptation phase
EMG: Surface electromyography
EW: Early washout phase
FAT: Fatigue group
LA_end_: Late adaptation phase – final block
LA_start_: Late adaptation phase – start
LB: Late baseline phase
LW: Late washout phase
MVC: Maximum voluntary contraction
NMF: neuromuscular fatigue
NoFAT: No-Fatigue group
RF: Rectus femoris
SOL: Soleus
TA: Tibialis anterior

## Notes

https://doi.org/10.17605/OSF.IO/A2XV4

### Competing Interest Statement

The authors have declared no competing interest.

### Summary of Updates

The current updated version implements revisions in: 1)Material and Methods section to improve clarity of data and statistical analyses; 2)Results section, implementing individual-level exponential fittings of performance error (PE) data, as well as detailed statistical outputs for additional postural measures; 3)Discussion section has been edited accordingly, to address study limitations. The Supplementary Data have been updated, adding a table (S6) detailing valid EMG data for each muscle used in the analyses.

