## Supplementary Data for "Adaptation to postural perturbations under fatigue produces persistent changes in neuromuscular coordination"

**Supplementary Materials**

**Figure S1:** Results for *CoP*$_{REC}$ *and Impulse*$_{AP}$ across the washout phase on each experimental session (*Day 1* and *Day 2*).

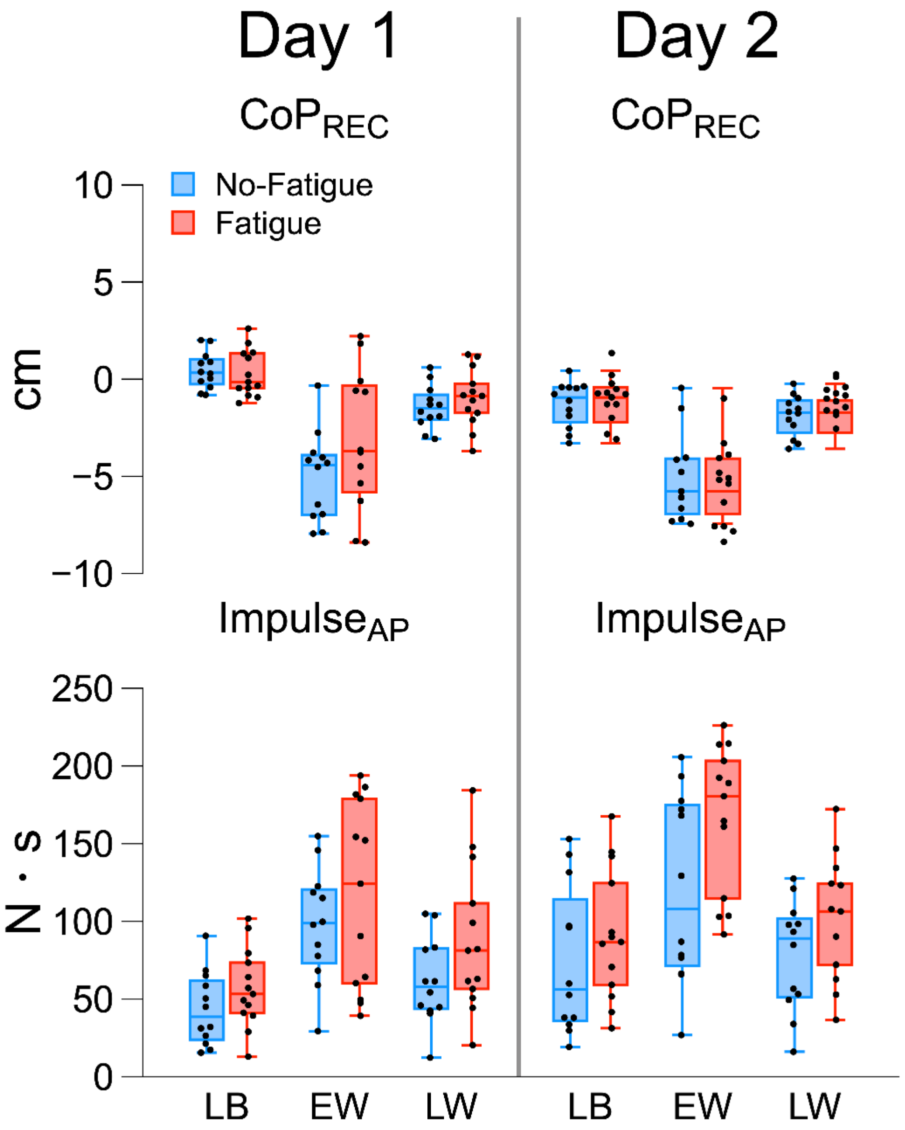


*Box plots for CoP*$_{REC}$ *and Impulse*$_{AP}$ *across the washout phase on each day. Boxes depict data from 1*$^{st}$ *to 3*$^{rd}$ *quartile; whiskers represent minimum and maximum data points. Median is shown by the thick horizontal line and individual data points as black dots. * Significant between/within group effects (*$p < 0.05$*). EW: early washout, LW: late washout, LB: late baseline.*

**Figure S2:** Results for EMGs of shank muscles (TA, SOL, *CC Index*$_{TA\text{-}SOL}$) across the washout phase on each experimental session (*Day 1* and *Day 2*).

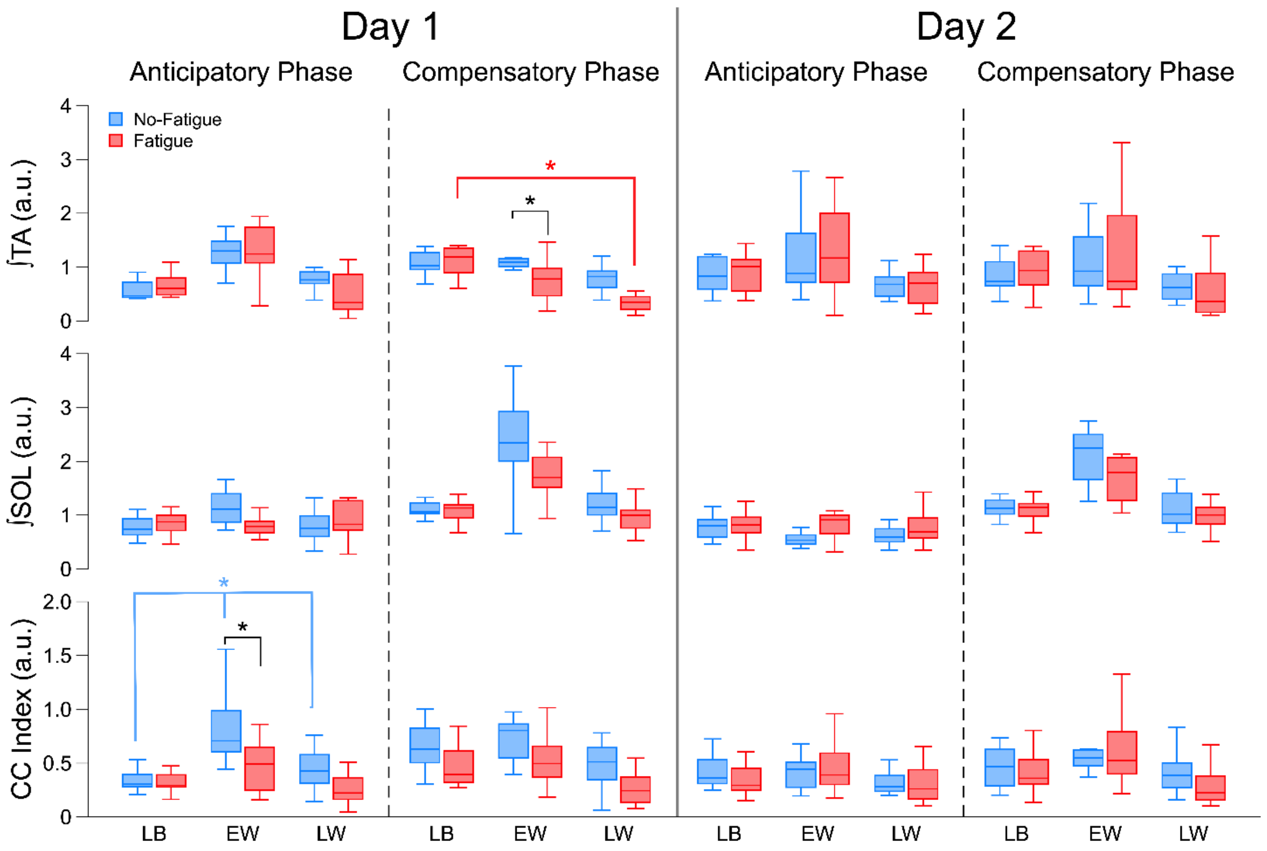


*Box plots with EMG activity of tibialis anterior (1st row), soleus (2nd row) and their coactivation index (CC Index; 3rd row) across the washout phase. The thick vertical line separates results by session (Day 1, Day 2), while pre-determined time window (anticipatory (AP) and compensatory (CP) phases) are shown side by side for each day. Boxes depict data from 1st to 3rd quartile; whiskers represent minimum and maximum data points. Median is shown by the thick horizontal line. * Significant between/within group effects ($p < 0.05$). EW: early washout, LW: late washout, LB: late baseline, SOL: soleus, TA: tibialis anterior.*

**Table S3:** ANOVA results for shank muscles EMG – *tibialis anterior* (TA) and *soleus* (SOL) – and their coactivation index (CC Index) in the washout phase. Results are reported separately for each session (*Day 1, Day 2*).

| Muscle | Anticipatory Phase (AP) | | | Compensatory Phase (CP) | | |
|---|---|---|---|---|---|---|
| | **Factor** | **F; p-value** | **Post-hoc** | **Factor** | **F; p-value** | **Post-hoc** |
| **Day 1** | | | | | | |
| **TA** | *Phase* | $F_{(2, 44)}$= 73.73, p<0.001, $\eta^2_p$ = 0.77 | All (p<0.001) | *Phase* | $F_{(2, 44)}$= 20.67, p<0.001, $\eta^2_p$ = 0.484 | |
| | *Group* | n.s | | *Group* | $F_{(1, 22)}$= 7.66 p=0.011, $\eta^2_p$ = 0.258 | EW (p=0.024) |
| | *Phase*Group* | n.s | | *Phase*Group* | $F_{(2, 44)}$= 4.04, p=0.025, $\eta^2_p$ = 0.155 | ***FAT only:*** LB–LW (p<0.001) |
| **SOL** | *Phase* | $F_{(2, 44)}$= 4.03, p=0.024, $\eta^2_p$ = 0.155 | n.s. | *Phase* | $F_{(1.23, 26.96)}$= 31.13, p<0.001, $\eta^2_p$ = 0.586 | LB–EW; EW–LW (p<0.001) |
| | *Group* | n.s. | | *Group* | n.s. | |
| | *Phase*Group* | $F_{(2, 44)}$= 3.53, p=0.038, $\eta^2_p$ = 0.138 | n.s. | *Phase*Group* | n.s. | |
| **CC Index** | *Phase* | $F_{(1.62, 34.01)}$= 16.87, p<0.001, $\eta^2_p$ = 0.446 | | *Phase* | $F_{(2, 42)}$= 14.13, p<0.001, $\eta^2_p$ = 0.402 | LB–LW; EW–LW (p<0.001) |
| | *Group* | $F_{(1, 21)}$= 14.3, p=0.001, $\eta^2_p$ = 0.406 | EW (p=0.015) | *Group* | $F_{(1, 21)}$= 8.48 p=0.008, $\eta^2_p$ = 0.288 | |
| | *Phase*Group* | $F_{(1.62, 34.01)}$= 3.75, p=0.042, $\eta^2_p$ = 0.152 | ***NoFAT only:*** LB–EW (p=0.002); EW–LW (p=0.016) | *Phase*Group* | n.s. | |
| **Day 2** | | | | | | |
| **Muscle** | Anticipatory Phase (AP) | | | Compensatory Phase (CP) | | |
| | **Factor** | **F; p-value** | **Post-hoc** | **Factor** | **F; p-value** | **Post-hoc** |
| **TA** | *Phase* | $F_{(1.18, 25.89)}$= 9.87, p<0.001, $\eta^2_p$ = 0.31 | LB–LW (p=0.011); EW–LW (p=0.004) | *Phase* | $F_{(1.14, 25.14)}$= 6.52, p=0.014, $\eta^2_p$ = 0.229 | LB–LW (p=0.043); EW–LW (p=0.012) |
| | *Group* | n.s | | *Group* | n.s | |
| | *Phase*Group* | n.s | | *Phase*Group* | n.s | |
| **SOL** | *Phase* | n.s | | *Phase* | $F_{(1.14, 25.14)}$= 27.38, p<0.001, $\eta^2_p$ = 0.554 | LB–EW; EW–LW (p<0.001) |
| | *Group* | n.s | | *Group* | n.s | |
| | *Phase*Group* | n.s | | *Phase*Group* | n.s | |
| **CC Index** | *Phase* | $F_{(1.31, 28.88)}$= 6.84, p=0.009, $\eta^2_p$ = 0.237 | LB–LW (p=0.007); EW–LW (p=0.01) | *Phase* | $F_{(1.42, 31.33)}$= 15.97, p<0.001, $\eta^2_p$ = 0.421 | All (p<0.05) |
| | *Group* | n.s | | *Group* | n.s | |
| | *Phase*Group* | n.s | | *Phase*Group* | n.s | |

Abbreviations: d: Cohen's d, $\eta^2_p$: partial eta squared, LB: late baseline, EW: early washout, LW: late washout.

**Figure S4:** Results for EMGs of thigh muscles (RF, BF, *CC Index$_{RF\text{-}BF}$*) across the washout phase on each experimental session (*Day 1* and *Day 2*).

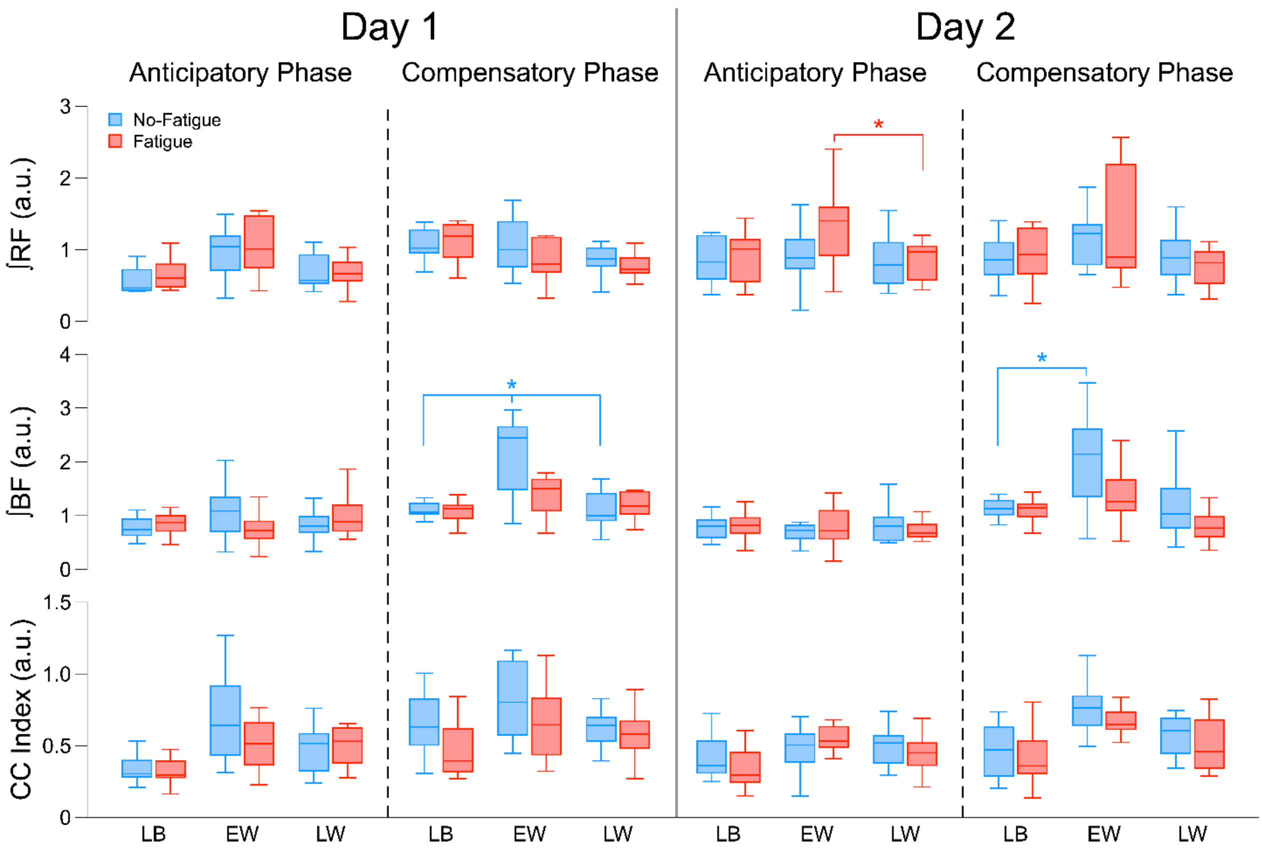


*Box plots with EMG activity of rectus femoris (1st row), biceps femoris (2nd row) and their coactivation index (CC Index; 3rd row) across the washout phase. The thick vertical line separates results by session (Day 1, Day 2), while pre-determined time window (anticipatory (AP) and compensatory (CP) phases) are shown side by side for each day. Boxes depict data from 1st to 3rd quartile; whiskers represent minimum and maximum data points. Median is shown by the thick horizontal line. * Significant between/within group effects ($p < 0.05$). BF: biceps femoris, EW: early washout, LW: late washout, LB: late baseline, RF: rectus femoris.*

**Table S5** ANOVA results for thigh muscles EMG – *rectus femoris* (RF) and *biceps femoris* (BF) – and their coactivation index (CC Index) in the washout phase. Results are reported separately for each session (*Day 1, Day 2*).

| **Day 1** | | | | | | |
|---|---|---|---|---|---|---|
| **Muscle** | *Anticipatory Phase (AP)* | | | *Compensatory Phase (CP)* | | |
| | **Factor** | **F; p-value;** | **Post-hoc** | **Factor** | **F; p-value** | **Post-hoc** |
| **RF** | *Phase* | $F_{(2, 44)}$= 7.08, p=0.002, $\eta^2_p$ = 0.244 | LB–EW (p=0.024); EW–LW (p=0.014) | *Phase* | n.s | |
| | *Group* | n.s | | *Group* | n.s | |
| | *Phase*Group* | n.s | | *Phase*Group* | n.s | |
| **BF** | *Phase* | n.s. | n.s. | *Phase* | $F_{(1.47, 32.3)}$= 20.34, p<0.001, $\eta^2_p$ = 0.48 | |
| | *Group* | n.s. | | *Group* | n.s. | |
| | *Phase*Group* | n.s. | | *Phase*Group* | $F_{(1.47, 32.3)}$= 4.79, p=0.024, $\eta^2_p$ = 0.179 | ***NoFAT only:*** LB–EW (p=0.002); EW–LW (p<0.001) |
| **CC Index** | *Phase* | n.s. | | *Phase* | $F_{(1.36, 28.49)}$= 4.18, p=0.039, $\eta^2_p$ = 0.166 | LB–LW (p=0.01) |
| | *Group* | n.s. | | *Group* | n.s. | |
| | *Phase*Group* | n.s. | | *Phase*Group* | n.s. | |

| **Day 2** | | | | | | |
|---|---|---|---|---|---|---|
| **Muscle** | *Anticipatory Phase (AP)* | | | *Compensatory Phase (CP)* | | |
| | **Factor** | **F; p-value** | **Post-hoc** | **Factor** | **F; p-value** | **Post-hoc** |
| **RF** | *Phase* | $F_{(1.44, 31.59)}$= 4.74, p=0.025, $\eta^2_p$ = 0.177 | ***FAT only:*** EW–LW (p=0.048) | *Phase* | n.s | |
| | *Group* | n.s | | *Group* | n.s | |
| | *Phase*Group* | n.s | | *Phase*Group* | n.s | |
| **BF** | *Phase* | n.s | | *Phase* | $F_{(2, 44)}$= 6.03, p<0.001, $\eta^2_p$ = 0.307 | ***NoFAT only:*** LB–EW (p=0.003) |
| | *Group* | n.s | | *Group* | $F_{(1, 22)}$= 7.15, p=0.014, $\eta^2_p$ = 0.245 | |
| | *Phase*Group* | n.s | | *Phase*Group* | n.s. | |
| **CC Index** | *Phase* | n.s | | *Phase* | $F_{(2, 44)}$= 7.25, p=0.002, $\eta^2_p$ = 0.248 | LB–LW (p=0.049); EW–LW (p=0.006) |
| | *Group* | n.s | | *Group* | n.s | |
| | *Phase*Group* | n.s | | *Phase*Group* | n.s | |

Abbreviations: d: Cohen's d, $\eta^2_p$: partial eta squared, LB: late baseline, EW: early washout, LW: late washout

**Table S6:** The table reports the number of participants included in each repeated measures ANOVA analysis for EMG measures, after listwise exclusion of invalid recordings. The starting sample was 25 participants.

| Measure | Day 1 AP | Day 1 CP | Day 2 AP | Day 2 CP |
|---|---|---|---|---|
| TA | 24 | 24 | 25 | 25 |
| SOL | 24 | 24 | 25 | 25 |
| CC Index$_{TA–SOL}$ | 22 | 23 | 25 | 25 |
| RF | 23 | 24 | 23 | 25 |
| BF | 24 | 23 | 25 | 25 |
| CC Index$_{RF–BF}$ | 23 | 22 | 25 | 25 |

**Note.** Reported values are the number of participants (N = 25 maximum). AP, anticipatory phase; CP, compensatory phase; CC Index, coactivation index; TA, tibialis anterior; SOL, soleus; RF, rectus femoris; BF, biceps femoris. Each ANOVA analysis used listwise deletion for any missing value across the experimental phases due to discarded invalid EMG recordings.
